# Age-dependent dynamics of neuronal VAPB^ALS^ inclusions in the adult brain

**DOI:** 10.1101/2024.02.12.579854

**Authors:** Aparna Thulasidharan, Lovleen Garg, Shweta Tendulkar, Girish S Ratnaparkhi

**Author notes:** Washington University School of Medicine, St. Louis, MO 63110, USA.

## Abstract

Amyotrophic Lateral Sclerosis (ALS) is a relentlessly progressive and fatal disease, caused by the degeneration of upper and lower motor neurons within the brain and spinal cord in the ageing human. The dying neurons contain cytoplasmic inclusions linked to the onset and progression of the disease. Here, we use a *Drosophila* model of *ALS8 (VAP^P58S^)* to understand the modulation of these inclusions in the ageing adult brain.

The adult *VAP^P58S^* fly shows progressive deterioration in motor function till its demise 25 days post-eclosion. The density of VAP^P58S^-positive brain inclusions is stable for 5-15 days of age. In contrast, adding a single copy of *VAP^WT^* to the *VAP^P58S^* animal leads to a large decrease in inclusion density with concomitant rescue of motor function and lifespan. ER stress, a contributing factor in disease, shows reduction with ageing for the disease model. Autophagy, rather than the Ubiquitin Proteasome system, is the dominant mechanism for aggregate clearance.

We explored the ability of *Drosophila* Valosin-containing protein (VCP/TER94), the *ALS14* locus, which is involved in cellular protein clearance, to regulate age-dependent aggregation. Contrary to expectation, *TER94* overexpression increased VAP^P58S^ punctae density, while its knockdown led to enhanced clearance. Expression of a dominant positive allele, *TER94^R152H^,* further stabilised VAP^P58S^ puncta, cementing roles for an *ALS8-ALS14* axis. Our results are explained by a mechanism where autophagy is modulated by *TER94* knockdown.

Our study sheds light on the complex regulatory events involved in the neuronal maintenance of ALS8 aggregates, suggesting a context-dependent switch between proteasomal and autophagy-based mechanisms as the larvae develop into an adult. A deeper understanding of the nucleation and clearance of the inclusions, which affect cellular stress and function, is essential for understanding the initiation and progression of ALS.

## INTRODUCTION

Amyotrophic Lateral Sclerosis (ALS) is a highly debilitating and fatal neurodegenerative disorder involving the lower motor neurons, brainstem, and spinal cord (Pasinelli and Brown 2006; Mitchell and Borasio 2007). The prognosis post-diagnosis is often bleak, with most patients succumbing to the disease within 2 to 5 years (Cleveland and Rothstein 2001; Boylan 2015). The mechanisms underlying the initiation and progression of ALS are unknown, with various hypotheses being tested in animal disease models (Van blitterswijk *et al.* 2012; Leblond *et al.* 2014; Renton *et al.* 2014). Drugs used for treatment include Riluzole (Doble 1996), Edaravone (Yoshino and Kimura 2006), Toferson (Miller *et al.* 2022) and Relyvrio (Paganoni *et al.* 2020). The drugs can moderately slow motor function decline and increase life expectancy by up to six months but cannot cure or arrest the disease. A better understanding of the nature of ALS pathology and its detrimental effect on the working of a healthy and functional nervous system could provide valuable insight into the disease.

ALS is sporadic, with incidences (<10%) of patients showing familial inheritance (Kurland and Mulder 1955; Boylan 2015). Many familial ALS (fALS) loci have been discovered, with ∼18 broadly classified as definitive loci (https://alsod.ac.uk/)(Abel *et al.* 2012). Functionally, these loci code for proteins/RNA with diverse functions, underscoring the complex aetiology of the disease. Mutations at the loci, in most cases, lead to aggregation of the translated protein, which is a common theme in neurodegenerative disorders (Ross and Poirier 2004). *VAPB* was identified in a Brazilian family as the eighth causative ALS locus (Nishimura *et al.* 2004). VAPB is a type -II Endoplasmic Reticulum (ER)-resident protein with three distinct domains: the Major Sperm Protein Domain (MSP), the Coiled-Coil Domain (CCD) and the Trans-Membrane Domain (TMD) (Skehel *et al.* 1995; Nishimura *et al.* 1999; Tsuda *et al.* 2008). The *VAPB* mutation identified by the Zatz group was a Proline to Serine substitution at the 56^th^ position (P56S) in the MSP domain (Nishimura *et al.* 2004). Regarding dysfunction, the VAPB^P56S^ protein is associated with ubiquitinated VAPB-positive cytoplasmic aggregates (Suzuki *et al.* 2009) on or abutting the ER. The mutation also affected the cleavage of the MSP domain, its secretion, and subsequent anterograde (Tsuda *et al.* 2008) signaling at the neuromuscular junction (NMJ). VAPB and its fly ortholog VAP33 (*CG5014*, VAP here onwards) have also been shown to have an expansive network of physical and genetic interactors, which is also affected by the mutation (Huttlin *et al.* 2015), which gave rise to the possibility of a causative oligogenic pathway. Previous studies from our laboratory have uncovered functional interactions of VAP with the TOR signaling pathway (Deivasigamani *et al.* 2014) and other orthologous ALS loci. A mechanistic understanding of these interactions has been uncovered for definitive ALS loci such as SOD1 (Chaplot *et al.* 2019) and TER94 (Tendulkar *et al.* 2022), as well as for finding the relationship between TOR signalling and VAP aggregation (Chaplot *et al.* 2019).

At the time of identification, VAP and its homologues were believed to have functions at the NMJ, lipid transport and microtubule-associated functionality (Skehel *et al.* 1995; Pennetta *et al.* 2002). VAPB function was expanded by discovering its wide-ranging physical interactors (Loewen and Levine 2005; Huttlin *et al.* 2015; Murphy and Levine 2016).

Intriguingly, in the past decade, VAPB has emerged as a protein involved in maintaining intracellular membrane:membrane contact sites (Zhao *et al.* 2018; James and Kehlenbach 2021)(Voeltz *et al.* 2024), with VAPB positioned on the cytoplasmic face of the ER, and associating with a variety of partner proteins on abutting cellular membranes, maintaining contact site architecture. VAPB appears to act as an influencer, directing the flux of its interaction partners and affecting cellular function.

In this study, we have attempted to understand the context of VAPB aggregation by modelling ALS8 in the *Drosophila melanogaster* adult brain in an age-dependent manner. Using a null-rescue disease model developed by the Tsuda laboratory (Moustaqim-Barrette *et al.* 2014) that phenocopies significant aspects of the clinical presentation of human ALS, we have developed quantitative systems to assess aggregation in the ageing animal. The age-dependent deterioration in motor function and short lifespan, seen in the *VAP^P58S^* disease model, is rescued by adding one copy of *VAP^WT^*. Intriguingly, this rescue correlates with a decrease of VAP^P58S^ punctae with age in the presence of one copy of *VAP^WT^* but not that of *VAP^P58S^*. Further, our study identifies *TER94*, the *Drosophila* ortholog of *VCP/ALS14*, as a critical modulator of VAP^P58S^ aggregation, highlighting autophagic mechanisms that presumably play a primary role in the clearance of VAP^P58S^ puncta in the adult brain.

## RESULTS

To study aggregation in the context of clinical manifestations of ALS, we adapted a genetic model system previously employed by the Tsuda lab (Moustaqim-Barrette *et al.* 2014; Tendulkar *et al.* 2022). Here, *Δ166*, a strongly hypomorphic *VAP* allele, is rescued by native, *VAP* genomic promoter-driven constructs (either *genomic(g) VAP^P58S^* or *VAP^WT^*). Both constructs rescue the larval/pupal lethality of *Δ166* (Tendulkar *et al.* 2022). Post-eclosion, the *ΔVAP;+;gVAP^WT^* animals show lifespan and motor function at par with wild-type animals. The *ΔVAP;+; gVAP^P58S^* lines, however, show a reduced lifespan (Suppl. Fig 1A) and a progressive decline in motor function, consistent with previous observations (Moustaqim-Barrette *et al.* 2014; Tendulkar *et al.* 2022). Adding a copy of *VAP^WT^* to the *VAP^P58S^* line was observed to rescue the lifespan and motor defects (Suppl. Fig. 1)(Moustaqim-Barrette *et al.* 2014; Tendulkar *et al.* 2022).

### Punctate localization of VAP is seen in the genomic *VAP^P58S^* line, with the puncta size, density and intensity scaling with the dosage of the mutant protein

In an earlier study, we used a rabbit antibody against the VAP CCD domain to mark VAP inclusions in fixed cells and larval brains (Chaplot *et al.* 2019). The *ΔVAP;+; gVAP^P58S^* larval brain also shows punctate localization of VAP (**Fig.1 C**, arrows), as opposed to the cytoplasmic staining shown by *ΔVAP;+; gVAP^WT^* and *CS* larval brains (Fig.1 A-B).

**Figure 1.**
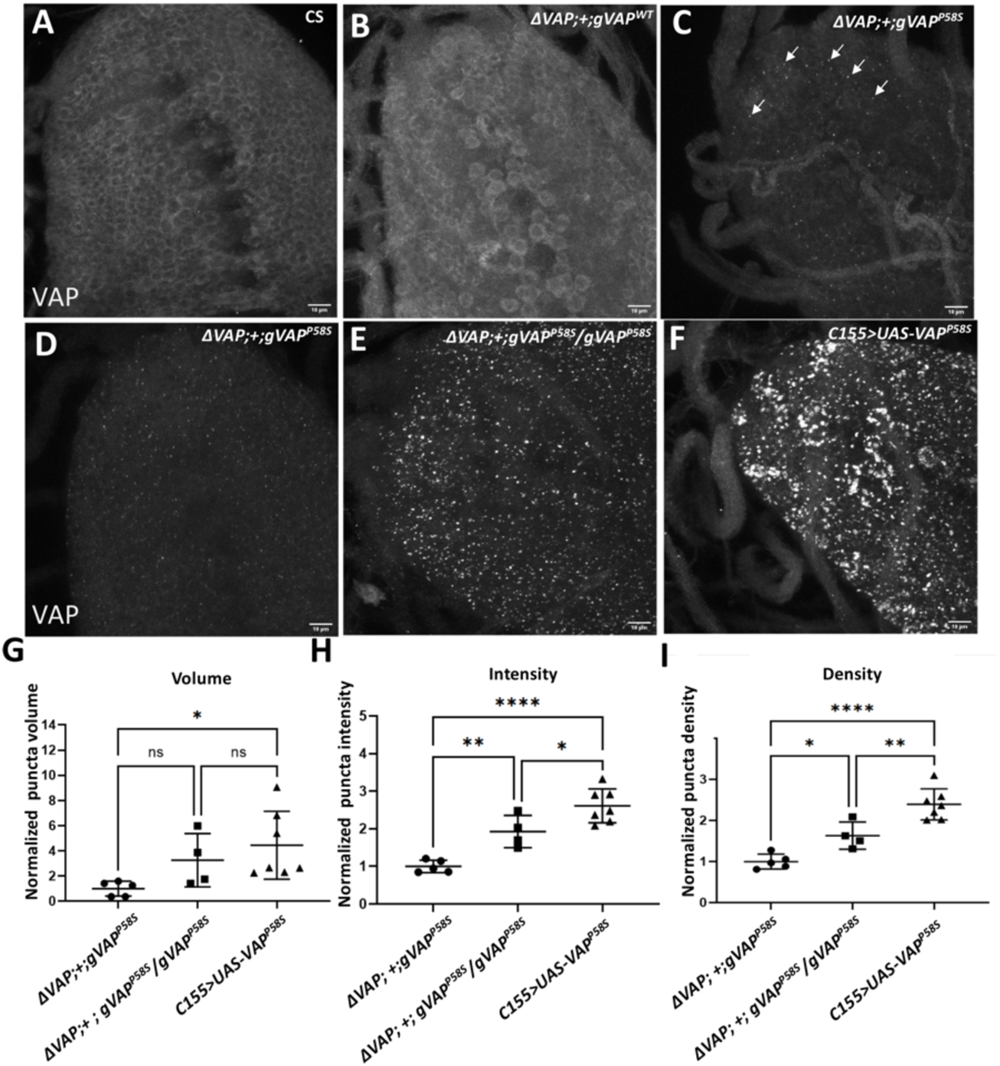
VAP inclusions show a dose-dependent increase in size, intensity and number. Maximum Intensity Projections (MIPs) of the larval VNCs of different genotypes stained with VAP antibody. Wild-type *Canton S, CS* (**A**) and *ΔVAP* rescued with g*VAP^WT^* (**B**) show non-punctate reticulate staining of VAP protein, marking the ER. *ΔVAP* rescued with a single copy of *VAP^P58S^* shows the presence of VAP positive puncta (white arrows, **C**). The puncta/inclusions increase in size, intensity and puncta density with the increase in *VAP^P58S^*dosage (**D-F**). The change in puncta volume (**G**), puncta intensity (**H**) and puncta density (**I**) over multiple experiments are depicted graphically. Each point **(G-I)** represents an average value calculated from 3 ROIs. For **G**, ns=non-significant,*p= 0.038. For **H**, *p= 0.033, **p=0.079,****p<0.0001. For **I**, *p=0.0287,**p= 0.0057, ****p<0.0001. The parametric test used is one-way ANOVA with Tukey’s multiple comparisons. Error bars represent SD. Scale bars denote 10µm.

We imaged larval brains (Fig. 1 D-F) with *VAP^P58S^* expression in a single genomic copy (Fig. 1 D), a double copy (Fig. 1 E), and overexpressed*-VAP^P58S^*, using the UAS-Gal4 system (Fig. 1 F). Next, we measured inclusion/puncta volumes, intensities and density at these varying dosages of *VAP^P58S^*. We observed an increase in puncta size, intensity and density with increased *VAP^P58S^* levels (Fig. 1 D-F, G-I). As before (Chaplot *et al.* 2019), we have used puncta density as the primary parameter to quantify changes in aggregation for the rest of this study.

After standardising our larval Ventral Nerve Cord (VNC) measurements, we expanded our study to include the adult brain. After staining with the anti-VAP antibody in the adult brain, we examined age-dependent (days 5, 11, 15) changes in aggregation for both a single *VAP^P58S^* and double dose *VAP^P58S/P58S^*of VAP. *ΔVAP;+;gVAP^WT^* animals (**Fig. 2 A-C**) did not show significant puncta, indicating absence of VAP aggregation. Both *ΔVAP;+;gVAP^P58S^* (Fig. 2 D-F) and *ΔVAP; gVAP^P58S^*/*gVAP^P58S^* (Fig. 2 G-I) animals showed VAP puncta. For the animals expressing VAP^P58S^ we did not observe a significant change in puncta density with age in either of the genotypes (Fig. 2 J). However, two copies of *gVAP^P58S^* showed a higher density of puncta formation, consistent with observations in the larvae (Fig. 2 J).

### *VAP^WT^* rescues aggregation in the larval ventral cord

One striking phenotype observed was the complete rescue of adult motor and lifespan defects observed when *VAP^WT^* was added to a *VAP^P58S^* animal. To study the molecular picture of this rescue, we compared puncta density in the larval VNCs between *ΔVAP;+;gVAP^WT^, ΔVAP;+; gVAP^P58S^*/*gVAP^WT^* and *ΔVAP; +; gVAP^P58S^* animals. The *ΔVAP; gVAP^WT^* animal showed ER staining (**Fig. 3 A**), while the *ΔVAP; +;gVAP^P58S^* animals have VAP positive puncta as the dominant feature (Fig. 3 B). In the *ΔVAP;+; gVAP^P58S^*/*gVAP^WT^* animals, we observe a significant decrease of puncta (Fig. 3 C) in the larval VNCs when compared to the *VAP^P58S^* animal. The VAP ER localization in the VNCs for *VAP^P58S^/VAP^WT^* larval brains closely resembles that of the wild-type alone (Fig. 3 A), with a small number of puncta and VAP present in the cytoplasm. This correlated well with the rescue of motor and lifespan defects, suggesting that VAP activity is normal in the *ΔVAP;+; gVAP^P58S^*/*gVAP^WT^* animals as a consequence of normal localization and function of VAP.

**Figure 2.**
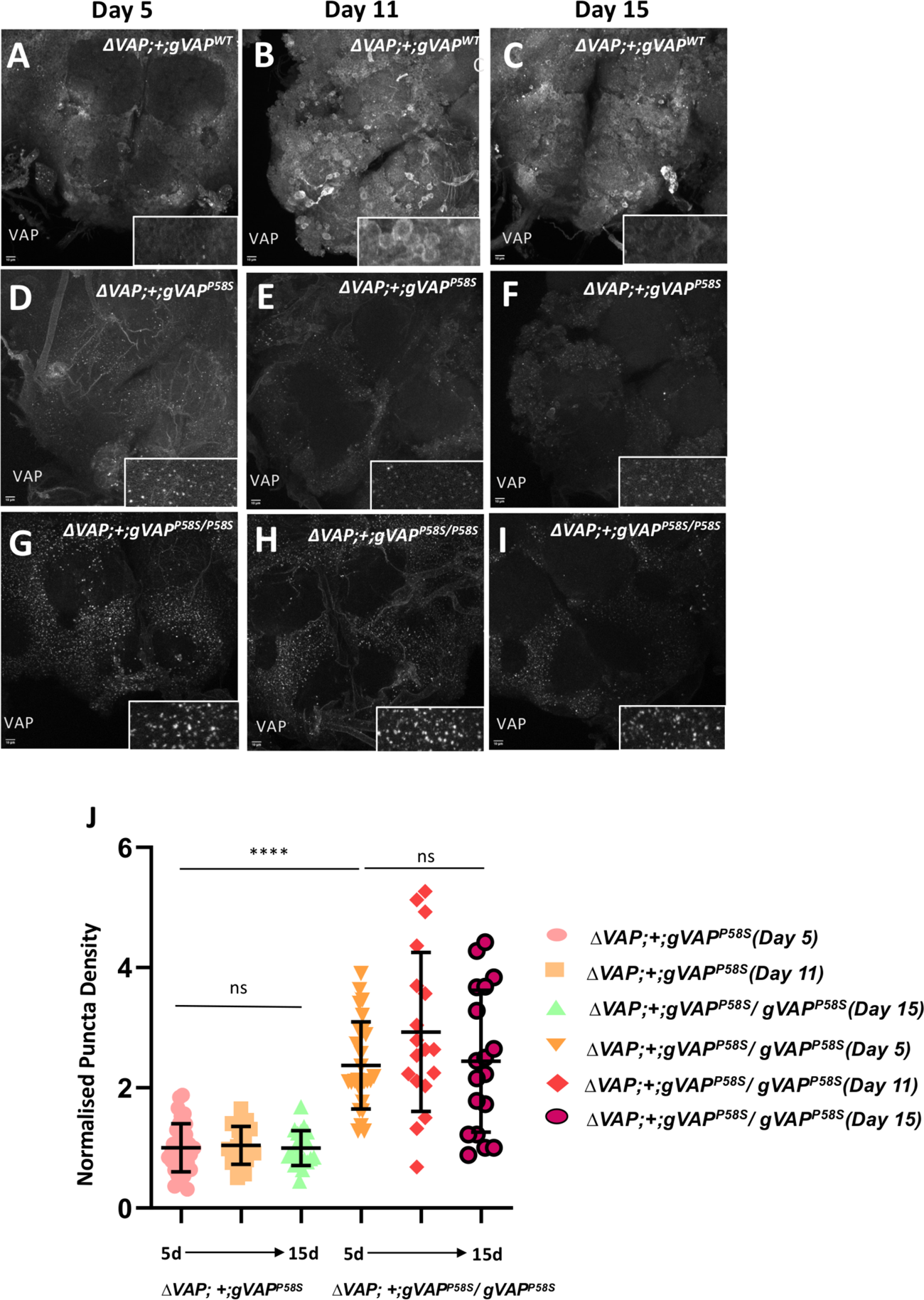
VAP puncta density does not change with age. Representative images of fixed adult brain anti-VAP staining for three-time points, day 5, day 11 and day 15. **(A-C)** *ΔVAP;+;gVAP^WT^*showing reticulate, non-punctate localization of VAP at all observed ages. **(D-F)** Δ*VAP;+;gVAP^P58S^*, and **(G-I)** *ΔVAP;+;gVAP^P58S^*/*gVAP^P58S^* show VAP positive puncta on all days. The punctae for *ΔVAP;+;gVAP^P58S^ ΔVAP;+;gVAP^P58S^*/*gVAP^P58S^*. Puncta density does not change across age **(J)** for each genotype, but the puncta density is higher when two copies of *gVAP^P58S^* are expressed, (G-I) as compared to one copy (D-F). ns=non-significant, ****P<0.0001. Kruskal-Wallis test, followed by Dunn’s multiple comparisons (n=3, N=8-12). Error bars depict SD). Scale bars denote 10µm.

**Figure 3.**
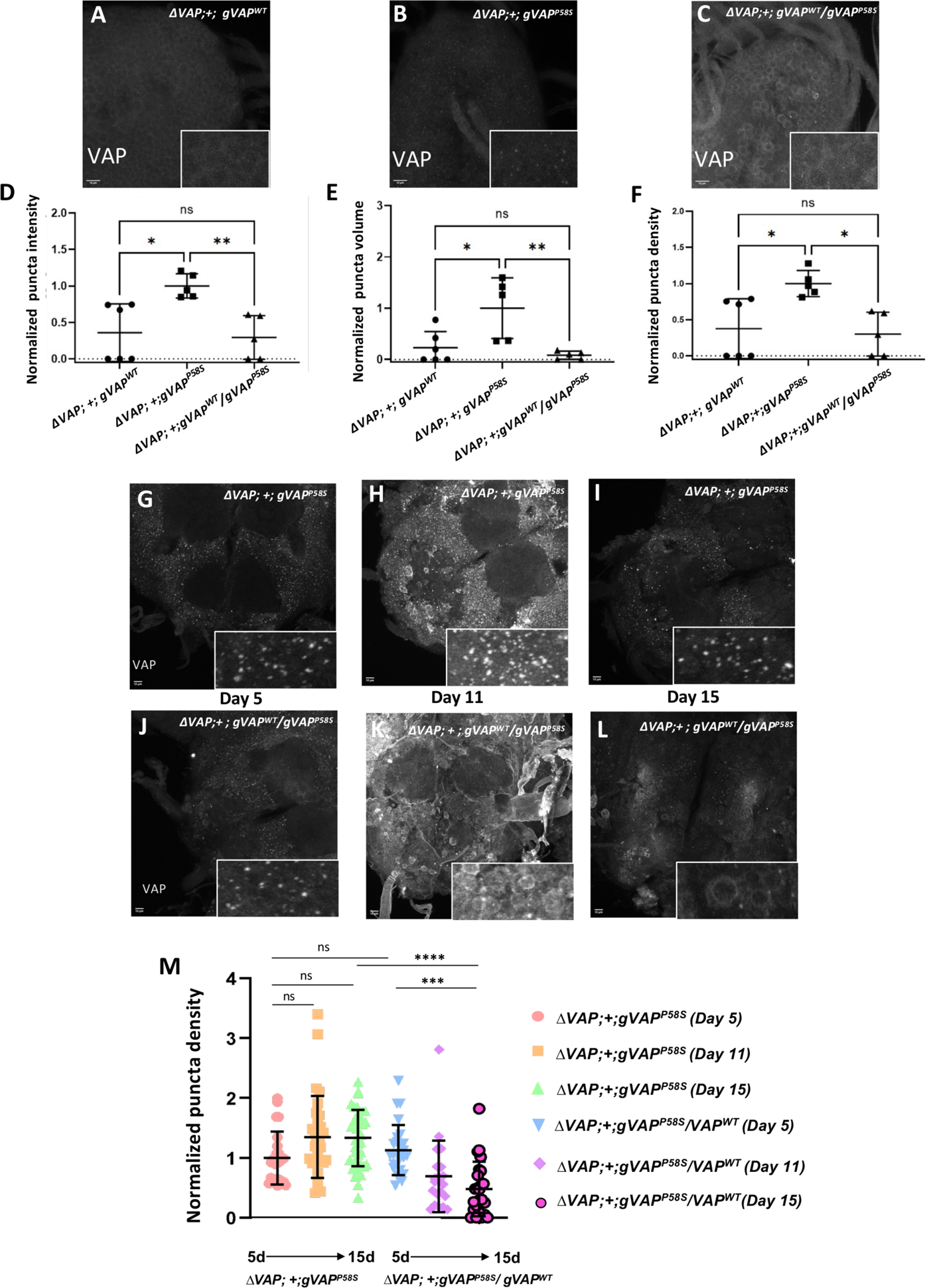
VAP puncta are cleared in the presence of a wild-type VAP allele. Figures (**A-C**) show representative Maximum Intensity Projections (MIPs) of the larval VNCs of *ΔVAP;+; gVAP^WT^, ΔVAP;+;gVAP^P58S^* and *ΔVAP;+;gVAP^P58S^ /gVAP^WT^* stained with VAP antibody. The VAP^P58S^ puncta intensity (**D**), volume (**E**) and puncta density (**F**) are reduced in the presence of VAP^WT^. Each point represents an average of 3 ROIs from one animal. (For **D**, ns=non-significant, *p= 0.012, **p= 0.0084. For **E**, *p=0.0144, **p=0.006. For **F**, *p= 0.01. One-way ANOVA with Tukey’s multiple comparisons was used for all three graphs. Error bars denote SD.) Scale bars denote 10µm. (**G-L**) Representative images of adult brain probed with anti-VAP antibody. *ΔVAP;+;gVAP^P58S^*and *ΔVAP;+ ;gVAP^P58S^/gVAP^WT^* across 5, 11 and 15 day old flies. Compared to *ΔVAP; +;gVAP^P58S^* brain (**G, H, I**), adults with a copy of *VAP^WT^* in the background **(J, K, L)** show a progressive decrease in puncta density. Quantifying punctae in terms of normalized puncta density, upon addition of *VAP^WT^* in the *VAP^P58S^* background (M). Each point represents a single ROI in a brain. *P=0.0187,***P=0.0002, ****P<0.0001, ns=Non significant, Kruskal-Wallis with Dunn’s multiple comparison (n=3, N=8-12). Error bars depict SD. Scale bar depicts 10µm.

We measured the puncta intensity, size and puncta density and the quantitative parameters measured agreed with our observations (Fig 3. D-F). In larvae, the presence of VAP^WT^ appeared to reduce the potential of VAP^P58S^ to aggregate. The next step was to look at the influence of VAP^WT^ on VAP^P58S^ inclusions in the adult brain.

### *VAP^WT^* reduces VAP^P58S^ aggregation in the adult brain in an age-dependent manner

In continuation with the experiments in the larval brain, we examined the brains of 5-day, 11-day and 15-day-old animals. As seen previously, the *ΔVAP;+; gVAP^P58S^* flies show puncta at all three age points (Fig. 3 G-I). There appears to be a slight increase in the average puncta density after day 5, but overall, the increase is insignificant (Fig. 3 G-I, M). However, in the case of *ΔVAP;+; gVAP^P58S^/gVAP^WT^* animals, the puncta density drops significantly with age, resulting in a clearance of the VAP puncta (Fig. 3 J-L, M). This indicates that the addition of *VAP^WT^* rescues VAP^P58S^ aggregation. The clearance is progressive, reducing with the age of the fly (Fig. 3 M). In the context of larval VNC clearance, it is interesting that even though larval brains have a low density of VAP^P58S^ aggregate in the presence of one copy of *VAP^WT^*, five days post-eclosion, the adult brain has similar normalized puncta density for VAP^P58S^ vs VAP^P58S^/VAP^WT^ (Fig. 3 M, 1 vs 1.2), when compared to VAP^P58S^/VAP^P58S^, which is ∼2.8 (Fig. 2 J).

Since aggregation can lead to ER stress, we next explored the effect of VAP^P58S^ aggregation and its clearance on ER stress over age.

### BiP aggregation reduces with age in the disease model; the presence of VAP^WT^ enhances this reduction

*VAPB* has previously been shown to be a regulator of the UPR and ER stress response (Suzuki *et al.* 2009). VAP null flies are known to show ER stress (Moustaqim-Barrette *et al.* 2014) and prolonged ER stress, which, if not managed and brought under control, can trigger pro-apoptotic pathways (Szegezdi *et al.* 2006). Overexpression of the mutant *VAP^P58S^* has also been shown to trigger ER stress and lead to BiP aggregation (Tsuda *et al.* 2008). BiP/Grp78 is an important player in the events leading to ER stress (Haas 1994). The binding of BiP to ATF6, PERK and IRE1 keeps them in an off-state. The presence of misfolded proteins causes BiP to dissociate from the complex and bind misfolded proteins instead, leading to Bip Puncta formation in the ER. BiP can attempt to refold the proteins by functioning as a chaperone (Lee 2005; Ibrahim *et al.* 2019). Once the misfolded proteins are removed, the BiP falls off the misfolded proteins and re-localises with ATF6, IRE1 and PERK (Bertolotti *et al.* 2000). As earlier studies were carried out in either a *VAP* null or with endogenous VAP in the background, we evaluated the age-dependent changes in Bip function in our disease model. We stained fly brains on day 15 to check for differences in BiP localisation between wild-type, *ΔVAP; gVAP^P58S^* and *ΔVAP; gVAP^P58S^/gVAP^WT^* (**Fig. 4**). We observed a very diffuse pattern of BiP staining in the case of 15-day-old wild-type flies indicating baseline levels of ER-stress.

**Figure 4.**
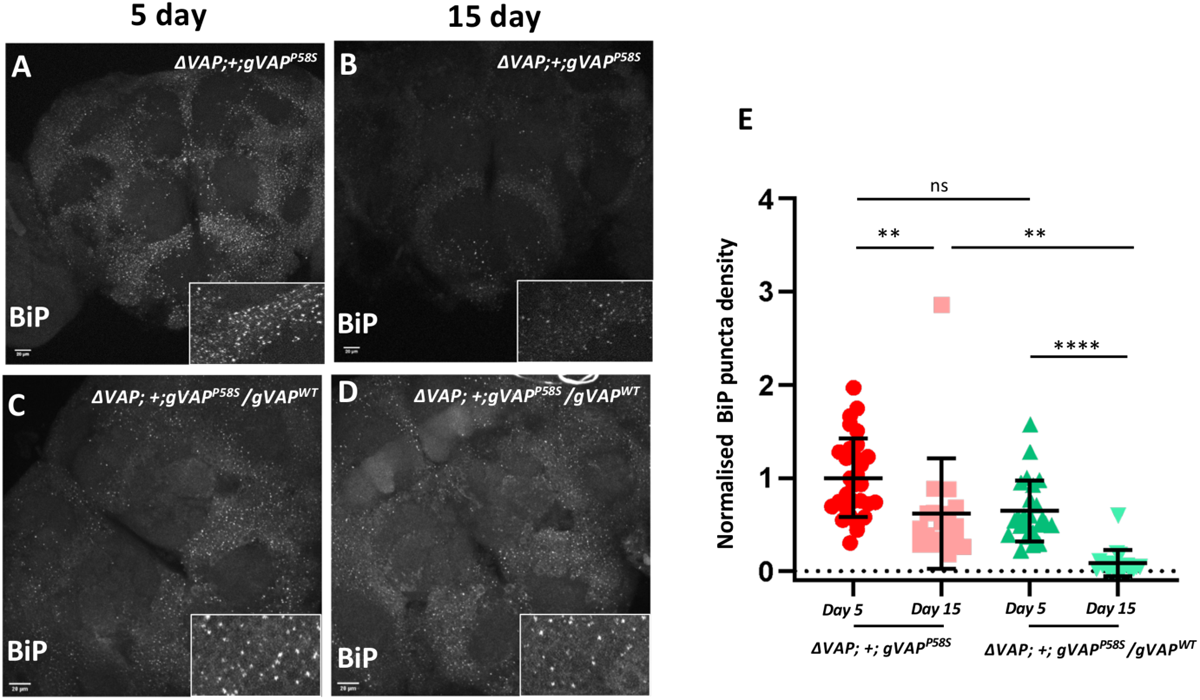
BiP aggregation reduces with age in the *VAP^P58S^* disease model; the presence of *VAP^WT^* enhances its reduction. BiP punctae were observed in *ΔVAP; gVAP^P58S^* (A and B), and *ΔVAP; gVAP^P58S^/gVAP^WT^* (C and D) animals at two time points, 5 and 15 days post eclosion. **(E)** BiP Aggregation density is lower at the 15 day time points for both genotypes. (A and B, **P=0.0074, for D and E, ****P<0.0001) when compared to 5 day animals. When comparing across genotypes at day 15 (B and D), **P=0.0034 and at day 5 (A and C), ns=non-significant. The test used was Kruskal-Wallis followed by Dunn’s multiple comparisons. Error bars depict SD. Scale bar represents 20 µm.

In contrast, the *ΔVAP; gVAP^P58S^* shows punctate BiP staining (Fig. 4 A), suggesting Bip binding to misfolded proteins. The puncta are indicative of compromised BiP activity and UPR. Comparison of the BiP puncta density of 5-day-old *ΔVAP; gVAP^P58S^* fly brains with 15-day-old *ΔVAP; gVAP^P58S^* fly brains (Fig. 4 A-B, E) shows a decrease in the puncta density at 15 days when compared to the 5-day old brains. The BiP aggregation phenotype does not appear to be progressive, instead it seems to ameliorate with age (Fig. 4 E). Interestingly, in the *ΔVAP; gVAP^P58S^/ gVAP^WT^,* we again observe a decline in BiP aggregation with age, but we see the aggregation drop to a far lower value of puncta density (Fig. 4 C-D, E). Thus, we find a reduction of BiP punctae with age in the disease model, suggesting a lowered ER stress response with an enhanced reduction in the presence of *VAP^WT^*.

### *SOD1* knockdown in the adult brain does not clear VAP^P58S^ aggregates

In the larval VNC, the knockdown of *SOD1* led to the proteasome’s clearance of the aggregated puncta in response to increased ROS levels (Chaplot *et al.* 2019). Surprisingly, in adult brains, we do not observe an equivalent phenomenon. We compared genotypes of *ΔVAP;Elav>+;gVAP^P58S^* and *ΔVAP; Elav>SOD1^RNAi^; gVAP^P58S^* fly lines in terms of puncta density (**Fig. 5 A-C**, ns, Mann-Whitney Test).

**Figure 5.**
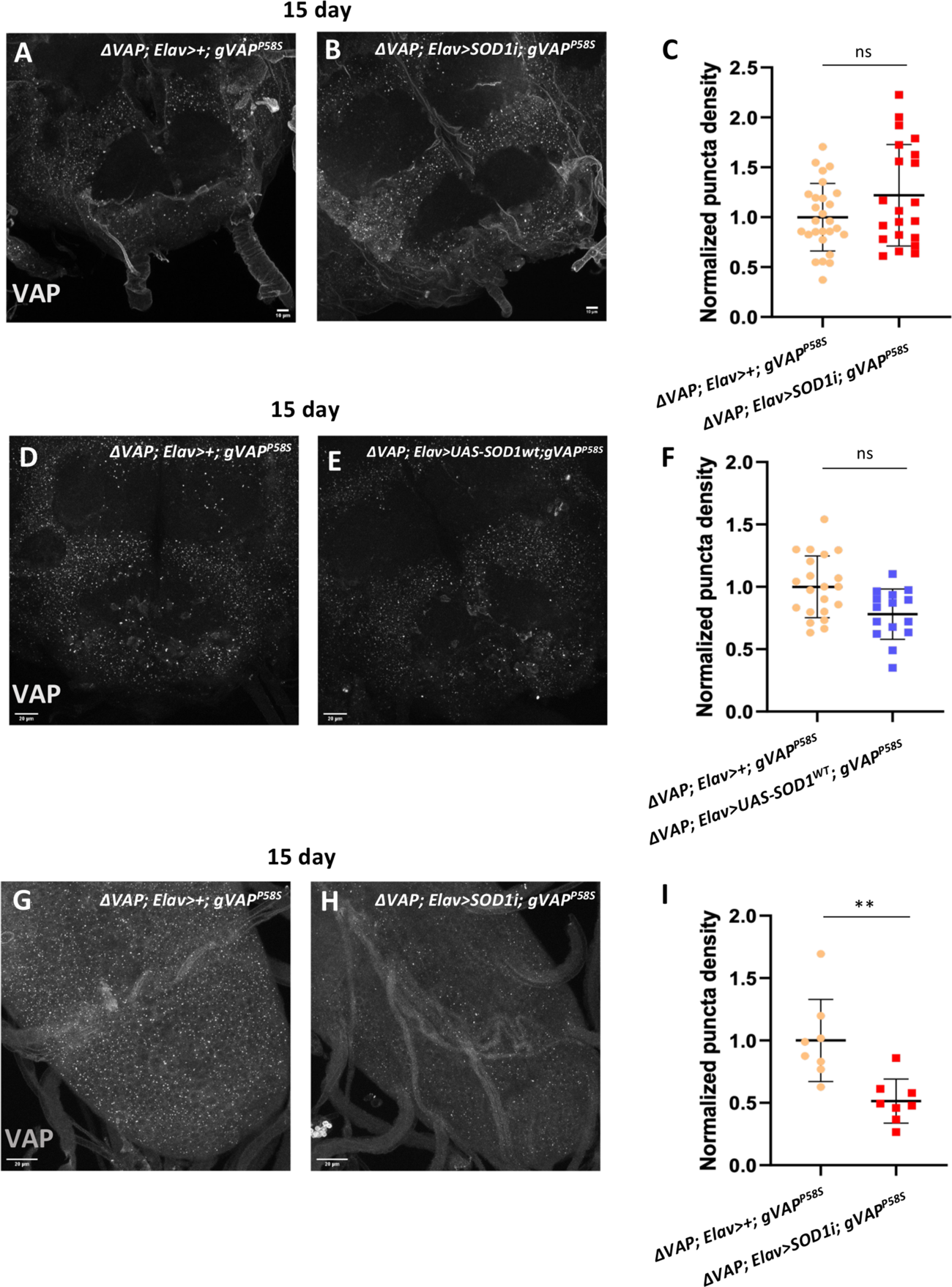
Knocking down *SOD1* neuronally does not change puncta density in the adult brain. Representative MIP images of the brains of *ΔVAP; Elav/+; gVAP^P58S^* **(A)** and *ΔVAP; Elav>SOD1i;gVAP^P58S^***(B)** stained with VAP antibody. The puncta density does not change between the two genotypes **(C)** (ns, Mann Whitney Test). Scale bar represents 10 µm **(D)** and **(E)** are representative MIP images of *ΔVAP;Elav/+;gVAP^P58S^ (*D) and *ΔVAP;Elav> UAS-SOD1 ^WT^;gVAP^P58S^*(E). The puncta density does not change between the two genotypes **(F)** (ns, Mann-Whitney test). Error bars represent SD. **(G)** and **(H)** are representative MIPs of larval Ventral nerve chords stained with VAP antibody. In **(H)** SOD1 is knocked down neuronally using the *Elav* driver. A reduction in VAP puncta density is observed when SOD1 is knocked down neuronally **(I)**. Each point represents an average value of puncta density derived from three ROIs per VNC. **P= 0.001, Mann-Whitney Test. Error bars represent SD. Scale bar denotes 20µm.

Here, the knockdown of *SOD1* did not affect puncta density, suggesting that ROS increase does not trigger proteasomal clearance in the adult brain. We also tried neuronally overexpressing *SOD1^WT^* in the *ΔVAP; Elav>+;gVAP^P58S^* animals (Fig. 5 D-F), which, too, did not appear to affect the puncta density significantly (Fig, 5 F, ns, Mann Whitney Test). Since the earlier larval experiments were conducted in an overexpression system (Chaplot *et al.* 2019), we repeated the larval VNC experiments afresh in the null-genomic rescue system. We examined *ΔVAP;Elav>+;gVAP^P58S^* and the *ΔVAP;Elav>SOD1^RNAi^;gVAP^P58S^* larval brains (Fig. 5 G-H) and calculated puncta density (Fig. 5 I). We saw a decrease in puncta density (Fig. 5I,**P=0.001, Mann-Whitney Test) in *SOD1i* knockdown. This result highlighted that ROS-mediated proteasomal clearance did not function in the adult brain as in the larvae.

The above observations suggest that the adult and larval mechanisms for proteostatic clearance of VAP aggregates differ and may constitute a context-dependent activation of different clearance pathways at various stages of the animal’s life cycle.

### TER94, the fly ortholog of VCP/p97 (ALS14), modulates VAP aggregation and the autophagic clearance of VAPB^P58S^ aggregates

The absence of proteasomal clearance in previous experiments led us to consider Autophagy as an alternate pathway for the clearance of VAP^P58S^ aggregates. This pathway was not dominant in the larval brain (Chaplot *et al.* 2019), where it was tested in the VAP overexpression model with VAP^WT^ in the background. To see if Autophagy was active in the adult brain, we knocked down *ATG1* using RNAi interference. ATG1 is a *Drosophila* ortholog of unc-51-like kinase 1 (ULK1) and an early and critical activator of the autophagic process (Mulakkal *et al.* 2014). When we knocked down *ATG1* in neurons, the density of VAP-positive puncta increased significantly (**Fig. 6B**). This suggested that clearance of aggregates via Autophagy is active in adult brain neurons. In this experiment, we used the brains of 15-day-old female animals, where, by 15 days, the presence of *VAP^WT^* had significantly reduced the density of VAP-positive puncta.

One long-term goal of our studies is to establish physiological relationships between orthologous ALS loci in the fly (Deivasigamani *et al.* 2014; Tendulkar *et al.* 2022). An earlier study (Tendulkar *et al.* 2022) established that *VCP/ALS14* is a significant genetic and physical interactor of *VAPB*. Both VCP (here onwards for flies, called *Drosophila* TER94, Transient Endoplasmic Reticulum 94 kD protein and VCP/p97 for mammals) and VAP physically interact with Caspar (Kim *et al.* 2006; Kaduskar *et al.* 2020; Tendulkar *et al.* 2022), which is the fly ortholog of human Fas-associated factor 1 (FAF1, (Chu *et al.* 1995; Ryu and Kim 2001; Ryu *et al.* 2003; Park *et al.* 2007; Baron *et al.* 2014)), suggesting roles for a VAPB:Caspar:TER94 complex in ALS. In the following sections, we test TER94 and then Caspar for roles in age-dependent aggregate dynamics.

**Figure 6.**
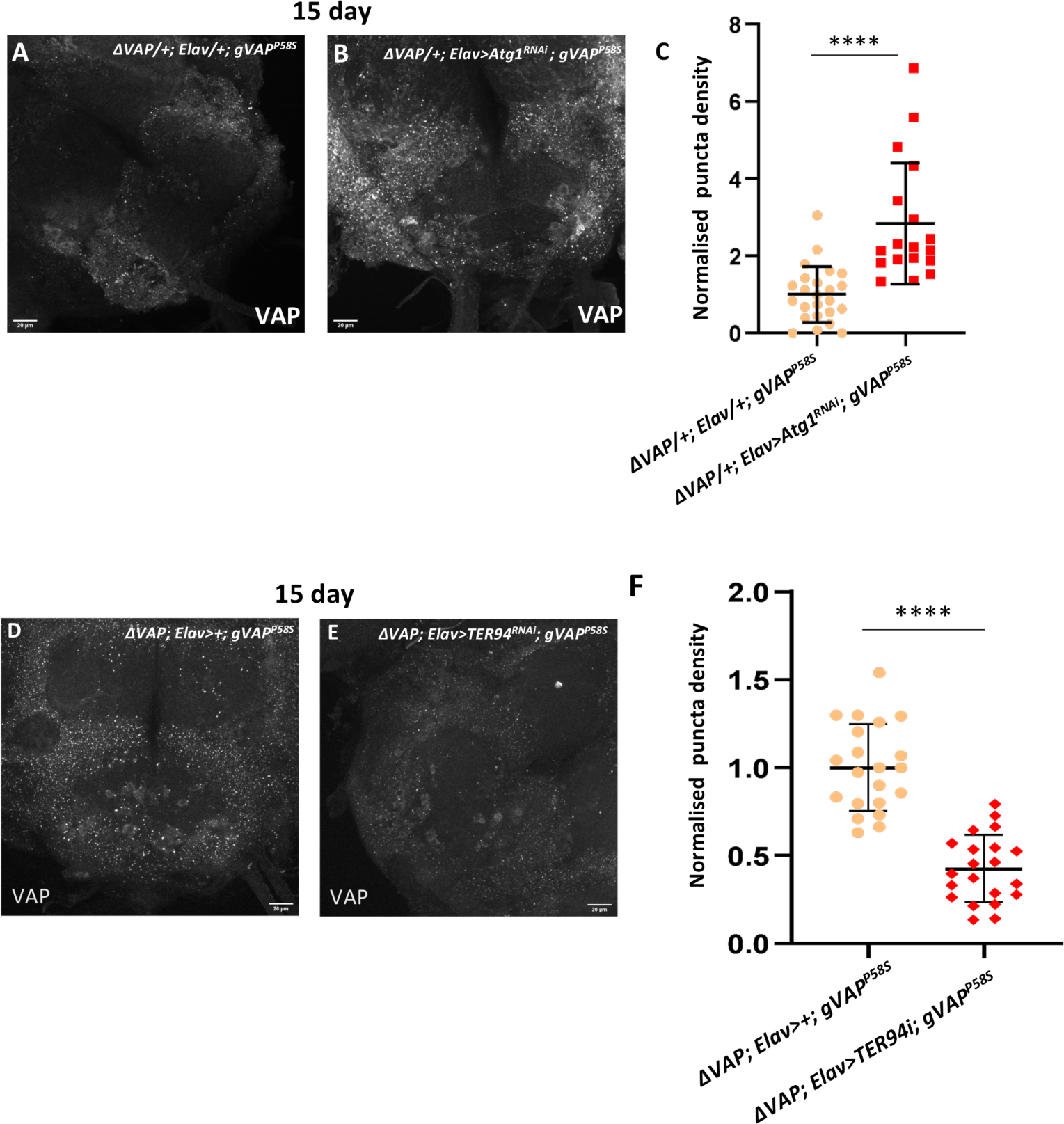
Autophagy and TER94 play roles in aggregate clearance/stability. Representative MIP images of the brains of 15 day old animals stained with anti-VAP antibody **(A)** *ΔVAP/+;Elav/+;gVAP^P58S^***(B)** *ΔVAP/+;Elav>Atg1^RNAi^;gVAP^P58S^*. There is an increase in puncta density upon neuronal knockdown of *Atg1* **(C)** *ΔVAP/+;Elav/+;gVAP^P58S^* and **(D)** *ΔVAP; Elav>Ter94^RNAi^;gVAP^P58S^*. The puncta density in the *TER94* neuronal knockdown is significantly reduced (C). ****P< 0.0001, Mann-Whitney Test. Error bars represent SD. Scale bar denotes 20µm.

TER94/VCP is a known component of the ERAD pathway and is involved in removing misfolded proteins (Leon and Mckearin 1999; Lee *et al.* 2013; Baron *et al.* 2014; Ahlstedt *et al.* 2022). VCP has also been implicated in Autophagy (Ju *et al.* 2008; Ju *et al.* 2009). In a paper from the Kundu laboratory, VCP was identified as a substrate for ULK1/2 (Wang *et al.* 2019). Phosphorylation of VCP by ULK1 increased the ATPase activity of VCP and led to efficient disassembly of stress granules (Wang *et al.* 2019). This suggested that *ATG1* knockdown may also reduce TER94 activity. Based on the above finding, reduction of TER94 activity is therefore predicted to reduce Autophagy dependent clearance, phenocopying the *ATG1* knockdown. To test this idea, *TER94* was knocked down in the neurons using the ELAV driver (*ΔVAP; Elav>Ter94^RNAi^; gVAP^P58S^*, Fig. 6E) and puncta density compared to control (*ΔVAP; Elav>+; gVAP^P58S^*, Fig. 6D). Surprisingly, we observed a sharp decrease in puncta density in the *TER94* neuronal knockdown (Fig. 6F). This result indicates that TER94 is indeed a player in the regulation of VAP aggregation. Still, the observations were not in line with our prediction.

### Increased TER94 activity stabilizes VAPB^P58S^ aggregates

In the previous section, intriguingly, the knockdown of TER94 activity reduced puncta density, suggesting clearance of aggregates. This is counterintuitive as TER94 has a canonical function as an ATP-dependent motor protein that assists in the clearance of proteins that require extraction from inclusions or membranes. To study the effect of increased TER94 activity in neurons, we overexpressed *TER94* using the ELAV driver. We see a mild increase (**Fig. 7 A,B,D**) in puncta density, suggesting that TER94 activity supports the stabilisation of aggregates.

**Figure 7.**
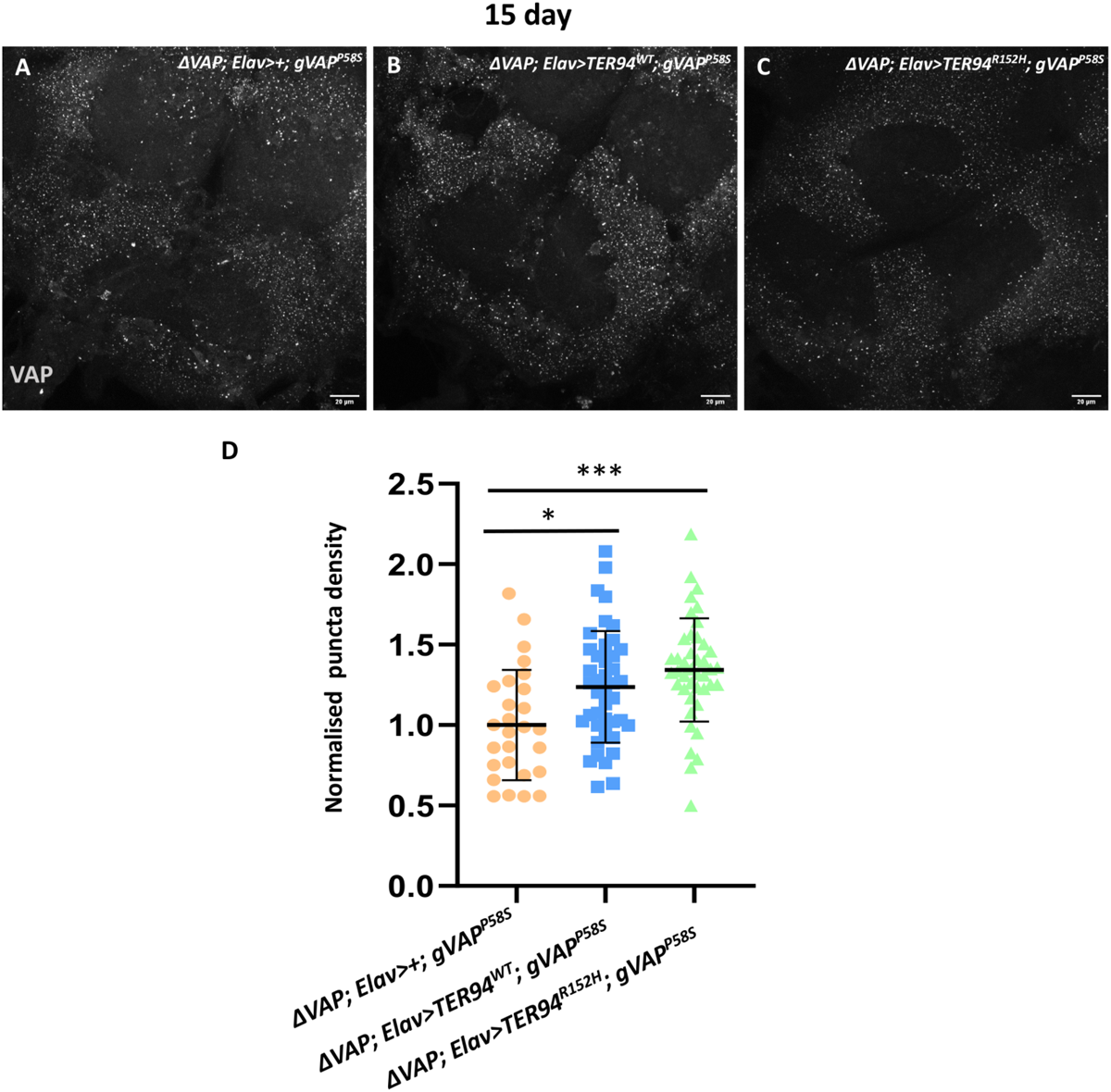
Neuronal overexpressing of TER94^R152H^ increases VAP aggregation density. Representative MIP images of the brains of 15 day old animals stained with anti-VAP antibody **(A)** *ΔVAP;Elav>TER94^WT^;gVAP^P58S^***(B)** *ΔVAP;Elav>+;gVAP^P58S^* and **(C)** *ΔVAP;Elav>TER94^R152H^;gVAP^P58S^*. The VAP puncta density for both the *TER94^WT^* and *TER94^R152H^* neuronal overexpression is higher **(D)** than that observed in the control. *P=0.0250,***P=0.0002, Kruskal-wallis followed by Dunn’s multiple comparisons. Error bars represent SD. The scale bar denotes 20µm.

The *VCP^R155H^* mutant is a known ALS-associated mutation (Johnson *et al.* 2010; Nalbandian *et al.* 2012), and the *TER94^R152H^* is the orthologous mutation in *Drosophila*. A *Drosophila* model developed for studying IBMPFD has defined *TER94^R152H^* as a dominant active allele (Chang *et al.* 2011). We observed a significantly higher density of VAP puncta when we overexpressed either *TER94^WT^* (Fig. 7B) or *TER94^R152H^* (Fig. 7 C-D, ***P=0.0002) as compared to the control (Fig. 7A). Thus, overexpression of both *TER94^WT^* and *TER94^R152H^* supported the formation or stabilisation of aggregates.

In summary, clearance mechanisms in the adult brain involved Autophagy, with TER94 modulating aggregate dynamics. Next, we tested if Caspar, a physical interactor of both VAP and TER94, had a role in maintaining VAP aggregates.

### Caspar does not modulate VAPB aggregation in neurons

Caspar (Casp) interacts with TER94, presumably via its UBX domain. Casp also interacts with VAP, via its FFAT-like motif (Tendulkar *et al.* 2022). This interaction is conserved in mammals, with VCP interacting with VAPB via its Casp ortholog, FAF1 (Baron *et al.* 2014). Many adapter proteins modulate the VCP/p97 function (Ewens *et al.* 2014; Zhang *et al.* 2015; Braxton and Southworth 2023). Adapter proteins contain VCP binding domains such as the UBX, UBXL, VIM, and VBM. The binding of these adapters can modulate the VCP function or give context to the VCP function (Chu *et al.* 2023). For example, the FAF1/p97 interaction promotes ERAD (Rabinovich *et al.* 2002; Lee *et al.* 2013). Previously, with studies focused on glia, we found that the knockdown of Casp in glia did not modulate puncta density but instead modulated the progression of the disease (Tendulkar *et al.* 2022), by regulating inflammation. We overexpressed Casp in neurons, *ΔVAP; Elav>Casp^WT^; gVAP^P58S^* (**Fig. 8 A**) and also knocked down using RNAi (Fig. 8 C; *ΔVAP; Elav>Casp^RNAi^; gVAP^P58S^*). The puncta density in 15-day adult brains was compared to controls (Fig. 8 B). No change was observed in puncta density (Fig. 8 D). This result indicates that Casp activity in neurons has no role in the context of VAP aggregate dynamics. Our results uncover an intriguing fact. In neurons, TER94 activity can modulate the stability of VAPB aggregates, but Casp, its binding partner and an interactor of VAP does not influence aggregation. In light of the potential roles for FAF1 in recruiting Ubiquinated proteins for degradation (Baron *et al.* 2014), this result of p97/VCP mediated proteasomal degradation further suggests a non-proteasomal route for VAP/VAPB degradation.

**Figure 8.**
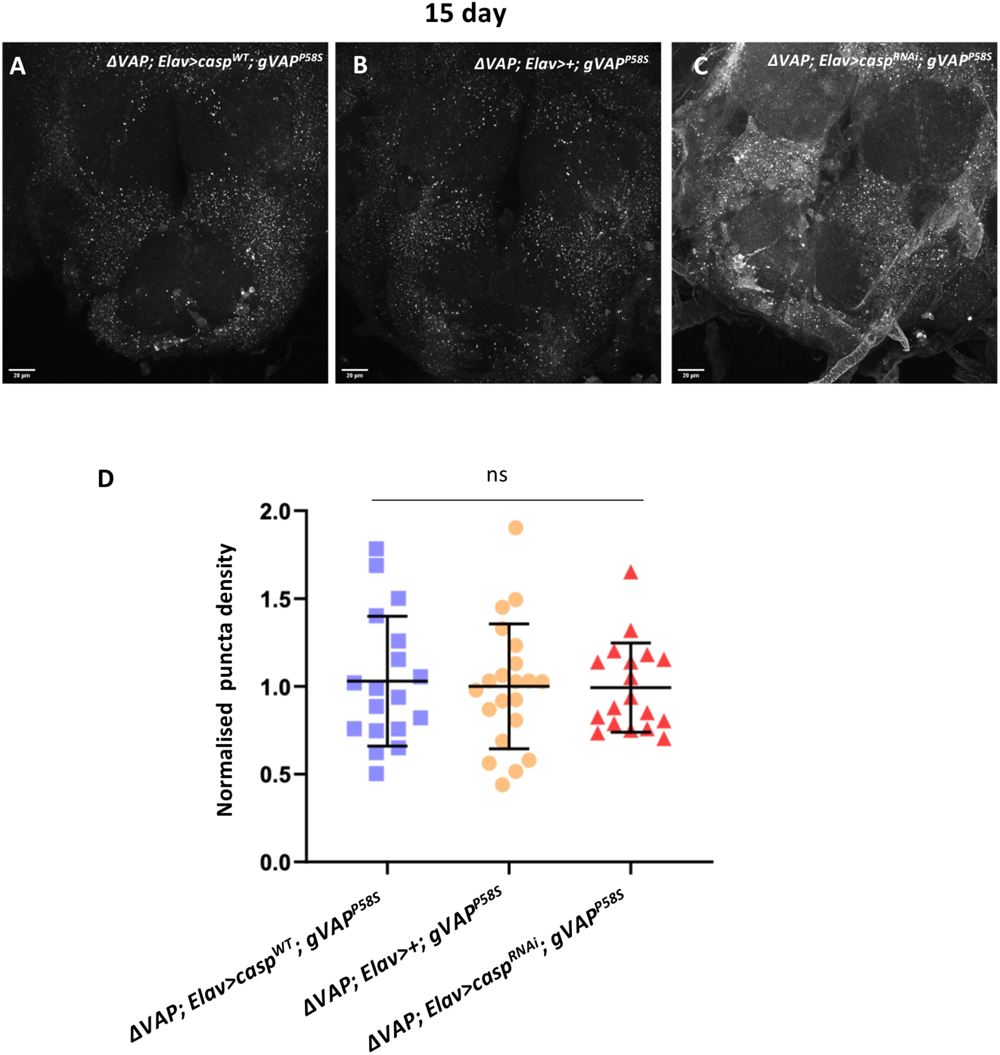
Modulation of *caspar* levels in the neurons does not affect VAP aggregation density. Representative MIP images of the brains of 15 day old animals stained with anti-VAP antibody **(A)** *ΔVAP; Elav>casp^WT^;gVAP^P58S^*, **(B)** *ΔVAP;Elav>+;gVAP^P58S^*and **(C)** *ΔVAP; Elav>casp^RNAi^;gVAP^P58S^* . The aggregation density **(D)** is similar for all three genotypes. Kruskal-Wallis followed by Dunn’s multiple comparisons. Error bars represent SD. The scale bar denotes 20µm.

## MATERIALS & METHODS

### *Drosophila* maintenance and husbandry

All flies were grown on standard corn meal agar at 25 °C, and all crosses were set up at 25 °C unless specified otherwise.

### *Drosophila* stocks and reagents

The flies expressing genomic(*g*) *VAP^WT^* and genomic *VAP^P58S^* were a gift from Hiroshi Tsuda (Moustaqim-Barrette *et al.* 2014). These lines were balanced with a strong, hypomorphic line, *Δ166*, as described previously (Tendulkar *et al.* 2022). Bloomington (BL) Stocks used in the study included BL 34616 (*UAS-SOD1^RNAi^*), BL 00001 (*CS*, *Cantos S*), BL 32869 (*UAS-VCP^RNAi^*) and BL 44034 (*UAS-ATG1^RNAi^*). CG11793-HA (*UAS-SOD1* overexpression) was procured from DPiM (https://interfly.med.harvard.edu/). *UAS-dVCP^WT^/CyO, UAS-dVCP^R152H^/CyO* are kind gifts from the Taylor laboratory (Ritson *et al.* 2010).

### Larval Ventral Nerve cord preparation

Wandering third-instar larvae were selected and dissected in 1X PBS. They were fixed in 4% PFA with 0.3% PBST for 20 minutes. Post fixation, the samples were washed thrice in 1X PBS and then transferred to blocking solution (2% BSA in 0.3% PBST) for 1 hour, followed by incubation in primary antibody for 1 overnight. This was followed by three washes with blocking each lasting 20 minutes and then an overnight incubation with the secondary antibody. Post-secondary, the samples are washed thrice with blocking, each lasting 20 minutes. DAPI is added to the second wash to visualise cell nuclei. Samples are given one final wash in PBS before mounting in antifade (Vectashield, S36937) mounting media.

### Adult brain dissections

Flies of the desired genotype are collected and aged to the requirement. Adult brains are dissected into cold 1X PBS, followed by a 24-hour fixation in 1.2% PFA at 4 °C. This is followed by permeabilization in 5% PBST (2 washes for 20 minutes each) followed by two washes in PAT buffer (PAT buffer is a 1X PBS buffer with 0.5% BSA and 0.5% Triton X), for 30 minutes each. Samples were blocked in 5% BSA in 0.5% PBST for two hours. This was followed by incubation in primary antibodies for 36 hours at 4 °C. Post-primary incubation, 4X PAT buffer washes, each lasting 30 minutes, were administered.

Samples are then incubated in secondary antibodies for 36 hours. Another four PAT buffer washes follow. DAPI (1:1000) is added in the second wash to visualize nuclei. This is followed by a wash in 1X PBS and subsequently stored in PBS with SlowFade:PBS in a 1:1 ratio at 4 °C overnight. For mounting, samples are placed with the antennal lobes facing the coverslip in a drop of Slowfade. Samples are bridge-mounted. Antibodies used are Rabbit Anti-VAP (1:500) (Yadav *et al.* 2018; Chaplot *et al.* 2019) and Rabbit anti-BiP (1:200)(Cell Signalling, C50B12) .

### Microscopy

Mounted samples were imaged using Zeiss LSM 710 or Leica SP8 confocal microscopes with 40X or 63X oil objectives. Images were acquired at 16-bit depth as Z stacks. For larval ventral chords, the tip of the ventral cord was imaged at 1X zoom. For the adult brains, the main body of the brain was imaged at 0.75X zoom. The system used was kept constant for each entire set of experiments. Acquisition parameters were kept constant across experimental sets. Due to the nature of the antibody and brain tissue, samples are imaged within five days of processing to avoid loss of fluorescence signal.

### Image analysis

We have used ImageJ and Huygens Professional software for image segmentation and analysis. The analysis protocol is similar to that used in (Chaplot *et al.* 2019) with modifications for analysing adult brain images. A threshold is determined per image per genotype to segment high-intensity punctae from the background signal from the diffuse staining of soluble VAP. The threshold is manually adjusted for each genotype to segment punctae from the highly variable background VAP intensity. Object filters were used to remove objects larger than 200 voxels and smaller than 8 voxels. The punctae were quantified per cubic micrometre of the larval ventral nerve chord or the adult brain, and this value is termed puncta density. Three 3D ROIs of adult brains and five larvae are selected from each sample from either the tip of the VNC (larval) or the sub-oesophageal zone (SEZ) (adult) and measured. The puncta density for each ROI was normalized to the mean value for the control group in each experiment. We have used 6 to 12 brains per genotype per experiment. ROI volume has been calculated using the range of the Z-stack of the image.

## DISCUSSION

*VAPB* has been a prominent locus (Nishimura *et al.* 2004; Mitne-Neto *et al.* 2007; Borgese *et al.* 2021a) for understanding ALS. The P56S mutation has been studied extensively (Kanekura *et al.* 2006; Teuling *et al.* 2007; Chai *et al.* 2008; Ratnaparkhi *et al.* 2008; Tsuda *et al.* 2008; Fasana *et al.* 2010; Kuijpers *et al.* 2013; Moustaqim-Barrette *et al.* 2014; Larroquette *et al.* 2015) in a variety of model systems and a significant amount of data collected on possible cellular mechanisms that lead to the disease. Features of cellular malfunction that come out of the vast array of research conducted include the formation of polymeric, ubiquitin-positive VAP aggregates in the cytoplasm (Teuling *et al.* 2007; Ratnaparkhi *et al.* 2008), progressive age-dependent motor and lifespan defects in animal models (Moustaqim-Barrette *et al.* 2014; Larroquette *et al.* 2015; Tendulkar *et al.* 2022), the partial functionality of the mutant protein (Moustaqim-Barrette *et al.* 2014), ER stress (Kuijpers *et al.* 2013; Moustaqim-Barrette *et al.* 2014), and non-cell autonomous roles for the MSP domain (Tsuda *et al.* 2008; Han *et al.* 2012). In the fly model, the *VAP^P58S^* mutation appears to be hypomorphic, rescuing the lethality of the *VAP^null^*. Further, adding a single copy of VAP^WT^ resolves lifespan and age-dependent, progressive motor defects that result from the VAP^P58S^ mutation (Moustaqim-Barrette *et al.* 2014).

### Ageing, Neuro-aggregates & Neurodegeneration

Ageing is a known risk factor for the development of several neurodegenerative disorders. Ageing leads to undesirable variations in cellular functions, including a decrease in proteostatic clearance, disruptions in autophagy, and increases in ROS and cellular inflammation. Studying neurodegeneration in the context of changes caused by ageing becomes essential for determining aetiology. While ageing has been shown to increase the concentration of insoluble proteins in cells (Reis-Rodrigues *et al.* 2012; Rai *et al.* 2021), and reduce proteasomal activity (Tsakiri *et al.* 2013), how it directly affects misfolding of proteins is not known. The presence of misfolded proteins that form large inclusions within and without the cell is a hallmark of neurodegenerative disease (Lim and Yue 2015; Wilson *et al.* 2023). In the case of ALS, the neuro aggregates are within the cell. The human VAPB^P56S^ aggregate has been modelled in multiple animal models, with inclusions seen in patient fibroblasts (Prause *et al.* 2013), cultured cell lines (Teuling *et al.* 2007; Fasana *et al.* 2010), mice (Larroquette *et al.* 2015), worms (Zhang *et al.* 2017) and flies. In flies, overexpression models have given way to systems where expression is regulated by native promoters (Moustaqim-Barrette *et al.* 2014). The VAP aggregates have been examined and well-characterized (Teuling *et al.* 2007; Mitne-Neto *et al.* 2011).

In our study, we utilized the Tsuda *Drosophila* model, which shows progressive motor defects and reduced lifespan, broadly in agreement with human ALS symptoms. Using this system, we could examine VAP puncta, a readout of VAP aggregation at different time points (ages) or genetic backgrounds. To study VAP aggregation and processes affected by it, we have developed a system for studying VAP protein aggregation using a workflow involving tissue dissection, immunostaining, fluorescence confocal microscopy and analysis of generated image stacks. In our study, we make critical observations regarding VAP aggregation with age. We observe that the puncta density does not vary significantly with the age of the fly. Motor defects progress with age, but such a correlation is not marked with increased VAP puncta formation, suggesting stable protein aggregation levels. This implies that the VAP punctae are not solely responsible for the progression of defects, and these may, in fact, not be toxic. The clearance of VAP^P58S^ aggregates in the presence of VAP^WT^ does, however, correlate with the animal’s lifespan and improved motor health.

Interestingly, ER stress, as measured by BiP positive puncta, goes down with age, even in the disease model, where motor function deteriorates with age and flies are destined to die by 20-25 days. The age-dependent decrease in ER stress seen in the disease model is enhanced with the expression of *VAP^WT^* in a *VAP^P58S^* animal. There is little doubt that VAPB^WT^ physiologically functions as a dimer or a higher-order multimer. Mutant and wild-type proteins associate together, as initially suggested (Ratnaparkhi *et al.* 2008). Our data indicates that the wild-type can stabilise the mutant, reduce aggregation/ puncta formation and support the animal for a full life span without apparent defects. Our data also supports the idea that in the case of ALS8, VAPB^P58S^ is partially functional, and haploinsufficiency is a significant contributor to the disease pathology. The concept of haploinsufficiency of VAPB is a familiar idea and has been discussed earlier (Borgese *et al.* 2021b). Thus, the decrease in cellular VAP is a significant factor in the disease and disease progression is assisted by ER stress (triggered by aggregates) and glial inflammation (Tendulkar *et al.* 2022). Lowered *VAPB* or VAPB or VAPB-activity levels are also features remarked on earlier in the case of patients (Mitne-Neto *et al.* 2011) or cultured cells (Papiani *et al.* 2012; Genevini *et al.* 2014). Further support comes from the fact that spinal cord lysates of ALS patients show low levels of VAPB (Tsuda *et al.* 2008).

### An unexpected role for TER94 in stabilising VAPB^P58S^ aggregates

The mechanism behind the clearance of VAP aggregates in larval brain knockdowns was the UPS (Chaplot *et al.* 2019), which was triggered by increased cellular ROS. This phenomenon appears not to be active in adult neurons. Instead, our data points to Autophagy being the dominant mechanism, with TER94 playing a significant role in autophagic clearance. TER94/VCP/p97 is an AAA-ATPase with canonical roles in extracting proteins for targeting the degradative pathway (Lord *et al.* 2002) (Meyer *et al.* 2012; Yamanaka *et al.* 2012; Baron *et al.* 2014; Van den boom and Meyer 2018). Proteins are extracted as part of ERAD (Wolf & Stolz, 2012, BBA)(Rabinovich *et al.* 2002) from membranes (Heo *et al.* 2010; Hemion *et al.* 2014) and large protein complexes or aggregates (Mukkavalli *et al.* 2021; Ahlstedt *et al.* 2022). Hexameric TER94 uses its central pore as a channel to extract unfolded proteins. For its function, it engages with numerous interacting factors that modulate TER94 activity. These factors provide context or specificity to the TER94 function and can enhance or block its activity.

A body of literature suggests that VCP or TER94 can disaggregate large protein inclusions (Mukkavalli *et al.* 2021). Our results, however, suggest the opposite; reduction in TER94 activity decreases the density of puncta-like aggregates while increasing activity stabilises aggregates. This phenomenon must be examined because Autophagy, rather than UPS, appears to be the predominant mechanism in clearing VAP aggregates in the adult brain. Past studies examining roles for VCP/p97 in autophagy predict that TER94 reduction would inhibit autophagy. For example, the Weihl laboratory (Ju *et al.* 2008; Ju *et al.* 2009) studying VCP in the context of inclusion body myopathy/Paget’s disease of the bone/frontotemporal dementia (FTD-IBMPFD) found that loss of VCP impairs autophagy.

Here, LC3-positive vacuoles fail to mature, accumulating nondegradative autophagosomes, leading to impaired degradation of protein aggregates. The Taylor lab (Tresse *et al.* 2010) independently confirmed this impairment. Further, Stress Granules also accumulate in conditions of VCP depletion (Seguin *et al.* 2014), underscoring VCPs positive role in aggregate clearance.

How do we explain our findings? Even though the larger body of work indicates that VCP/p97 inhibition reduces Autophagy, there are exceptions. CB-5083, an inhibitor of VCP/p97, enhances autophagy (Anderson *et al.* 2015) rather than impairs it, indicating that the phenomenon may be context-dependent. Another possibility is that under conditions of low TER94 activity, the clearance mechanisms may shift to Proteasomal degradation, a possibility not tested in our experiments. VAPB is a single pass, tail-anchored membrane protein and such proteins are inserted in the ER by a Guided entry of tail-anchored proteins (GET; (Borgese and Fasana 2011; Colombo and Fasana 2011; Hegde and Keenan 2011)). TER94 may play a non-canonical or undiscovered role in either stabilizing membrane insertion or in destabilising large cytoplasmic aggregates, which we visualise as puncta, thus influencing aggregate density.

Our overexpression data is more consistent with the literature. The *VCP^R155H^* allele is defined as a dominant active allele, and its expression leads to an increase in ubiquitin-conjugated proteins and the stabilisation of ΔF508-CFTR aggregates (Weihl *et al.* 2006).

*VCP^R155H^* impairs autophagosome maturation and fusion with lysosome (Ju *et al.* 2008; Ju *et al.* 2009; Tresse *et al.* 2010), blocking autophagic processes. Increased TER94 activity promotes the formation of Rhodopsin (Rh1) oligomers (Griciuc *et al.* 2010), broadly supporting a role for the generation of large aggregates. The *VCP^R155H^* has also been reported to increase the seeding of α-synuclein and TDP43 aggregates (Zhu *et al.* 2022). Also to be considered is the link of *VCP^R155H^* with the disease. (Griciuc *et al.* 2010) found that decreased *TER94* activity reduced retinal neurodegeneration. In contrast, the Taylor lab (Ritson *et al.* 2010), studying an IBMPFD model in the context of TDP43, found that exogenous expression of disease-related VCP mutations enhanced the toxicity of TDP43.

In summary (**Fig. 9**), a single missense mutation VAP^P58S^ leads to the formation of large cytoplasmic aggregates, with a significant fraction inserted in the ER (Fig. 9 A). As reported earlier (Papiani *et al.* 2012), VAPB^P56S^ can modify ER architecture. When on the ER, the mutant proteins function as hypomorphic alleles, providing wild-type function. On translation of *VAP*, large clusters of non-toxic VAP^P58S^ aggregates are accumulated in clusters akin to stress granules (Fig. 9 B), and these are visualised as puncta in our anti-VAP neuronal staining. These aggregates trigger an ER stress response. The addition of VAP^WT^ leads to the formation of heterodimeric VAP^WT^:VAP^P58S^. These hetero-multimers are less aggregation-prone and functionally at par with VAP^WT^: VAP^WT^. The VAP^WT^ stabilizes VAP^P58S^ (on ER, Fig. 9 C) and reduces puncta (Fig. 9 B) formation. Intriguingly, the clearance progresses with age.

Autophagy appears to be a dominant mode of clearance of puncta (Figure 9 D) and may also involve ER-phagy. The overexpression of *TER94R^152H^*, a disease ortholog, and a dominant active variant leads to increased puncta formation and may increase clusters on the ER (Fig. 9E). Despite our findings and those of earlier researchers, many open questions remain. We have much to learn about age-dependent aggregation, clearance mechanisms, and the role of VCP in Autophagy vs Proteasomal clearance.

**Figure 9.**
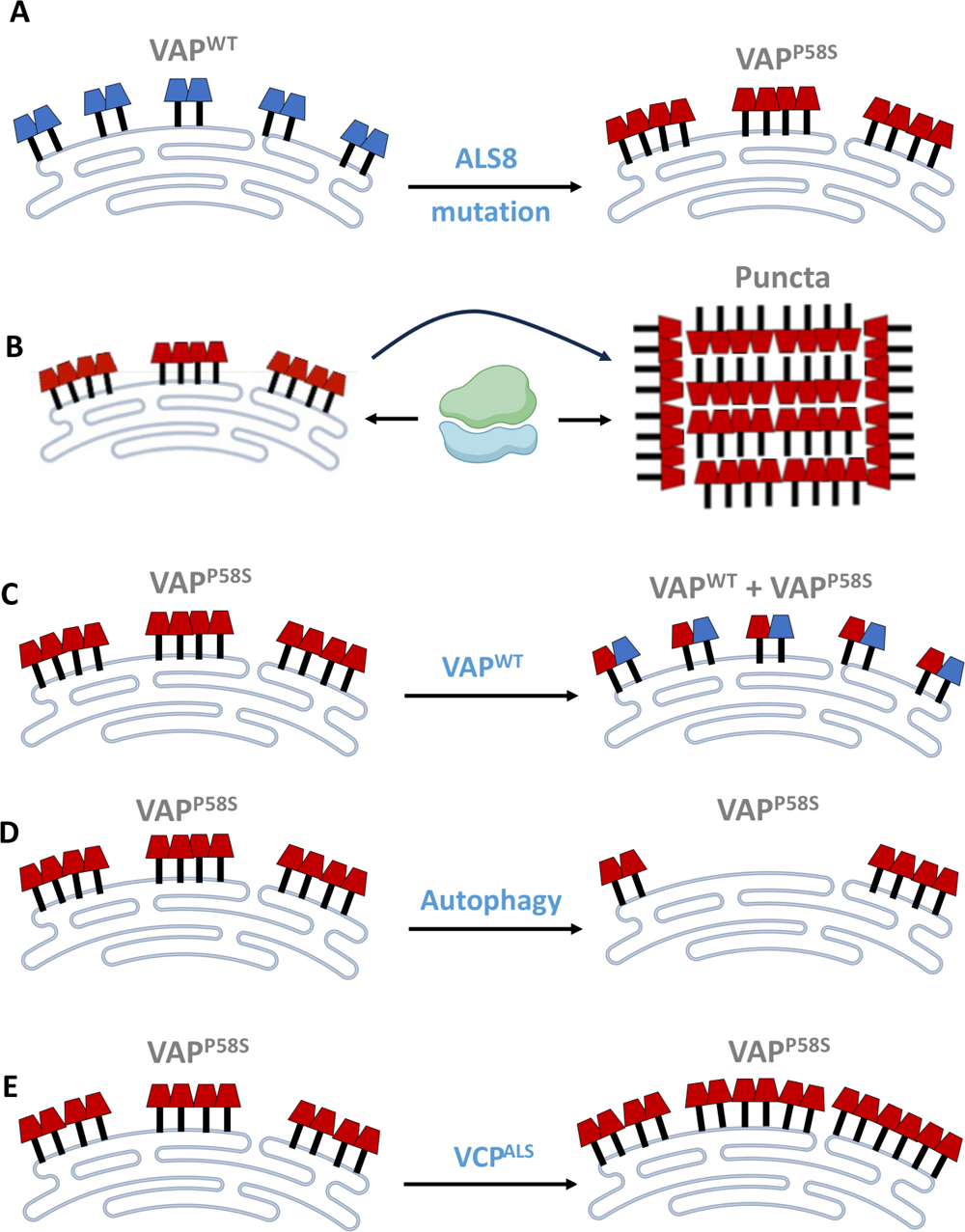
A model for the dynamics of VAPB^P58S^ aggregation and clearance. Our experiments, conducted in the *Drosophila* larvae and adult brain, suggest the following mechanisms for stabilisation and/or clearance of aggregates. **(A)** VAPB^WT^ protein (in blue) forms multimeric (dimeric/tetrameric) associations on the ER membrane, with the MSP domain facing the cytoplasm. The ALS-orthologous mutation (VAPB^P58S^) leads to large multimeric aggregates (red, clustered) that can be visualized as VAPB-positive puncta on antibody staining. **(B)** A significant fraction of the clusters will be ER-membrane-associated, as VAPB^P58S^ variant can support animal development **(C)** VAPB^WT^ protein facilitates an age-dependent decrease of puncta in the adult brain, suggesting clearance of VAPB^P58S^ puncta, and by forming heterodimers (VAPB^WT^:VAPB^P58S^) that reduce the size and incidence of cytoplasmic aggregates. The heterodimer appears at par with VAPB^WT^ in terms of function **(D)** Autophagy appears to be the dominant mechanism for the clearance of VAPB^P58S^ aggregates, with a reduction in autophagic clearance, by blocking *ATG1,* stabilizing aggregates and possibly reducing ER clusters (**E)** TER94 activity modulates VAPB^P58S^ aggregate formation with increasing activity (TER94^WT^ or TER94^R152H^) stabilizing aggregation, increasing puncta density. Reduced TER94 activity leads to the clearance of aggregates and reduced puncta density.

## CONTRIBUTIONS

AT and GR conceptualised the project and designed the experiments. AT performed all the experiments. LG was involved in data analysis, while ST balanced and characterised the disease model. AT and GR analyzed the data and wrote the manuscript. GR supervised the project and acquired funding.

## ACKNOWLEDGEMENTS

We thank Bloomington Drosophila Stock Center (BDSC), Indiana, supported by NIH grant P40OD018537, for fly stocks; JP Taylor for UAS-VAP lines; Snehal Patil and Yashwant Pawar for fly media and stock maintenance; IISER Microscopy facility, Dr Santosh Podder and Vijay Vittal for training and maintenance.

## REFERENCES

Abel, O., J. F. Powell, P. M. Andersen and A. Al-Chalabi, 2012 ALSoD: A user-friendly online bioinformatics tool for amyotrophic lateral sclerosis genetics. Human Mutation 33: 1345–1351.

Ahlstedt, B. A., R. Ganji and M. Raman, 2022 The functional importance of VCP to maintaining cellular protein homeostasis. Biochem Soc Trans 50: 1457–1469.

Anderson, D. J., R. Le Moigne, S. Djakovic, B. Kumar, J. Rice et al., 2015 Targeting the AAA ATPase p97 as an Approach to Treat Cancer through Disruption of Protein Homeostasis. Cancer Cell 28: 653–665.

Baron, Y., P. G. Pedrioli, K. Tyagi, C. Johnson, N. T. Wood et al., 2014 VAPB/ALS8 interacts with FFAT-like proteins including the p97 cofactor FAF1 and the ASNA1 ATPase. BMC Biol 12: 39.

Bertolotti, A., Y. Zhang, L. M. Hendershot, H. P. Harding and D. Ron, 2000 Dynamic interaction of BiP and ER stress transducers in the unfolded-protein response. Nat Cell Biol 2: 326–332.

Borgese, N., and E. Fasana, 2011 Targeting pathways of C-tail-anchored proteins. Biochim Biophys Acta 1808: 937–946.

Borgese, N., N. Iacomino, S. F. Colombo and F. Navone, 2021a The Link between VAPB Loss of Function and Amyotrophic Lateral Sclerosis. Cells 10: 1865.

Borgese, N., F. Navone, N. Nukina and T. Yamanaka, 2021b Mutant VAPB: Culprit or Innocent Bystander of Amyotrophic Lateral Sclerosis? Contact (Thousand Oaks) 4: 25152564211022515.

Boylan, K., 2015 Familial Amyotrophic Lateral Sclerosis. Neurol Clin 33: 807–830.

Braxton, J. R., and D. R. Southworth, 2023 Structural insights of the p97/VCP AAA+ ATPase: How adapter interactions coordinate diverse cellular functionality. J Biol Chem 299: 105182.

Chai, A., J. Withers, Y. H. Koh, K. Parry, H. Bao et al., 2008 hVAPB, the causative gene of a heterogeneous group of motor neuron diseases in humans, is functionally interchangeable with its Drosophila homologue DVAP-33A at the neuromuscular junction. Hum Mol Genet 17: 266–280.

Chang, Y. C., W. T. Hung, Y. C. Chang, H. C. Chang, C. L. Wu et al., 2011 Pathogenic VCP/TER94 Alleles Are Dominant Actives and Contribute to Neurodegeneration by Altering Cellular ATP Level in a Drosophila IBMPFD Model. Plos Genetics 7: e1001288.

Chaplot, K., L. Pimpale, B. Ramalingam, S. Deivasigamani, S. S. Kamat et al., 2019 SOD1 activity threshold and TOR signalling modulate VAP(P58S) aggregation via reactive oxygen species-induced proteasomal degradation in a Drosophila model of amyotrophic lateral sclerosis. Disease Models & Mechanisms 12: dmm033803.

Chu, K. T., X. H. Niu and L. T. Williams, 1995 A Fas-Associated Protein Factor, Faf1, Potentiates Fas-Mediated Apoptosis. Proceedings of the National Academy of Sciences of the United States of America 92: 11894–11898.

Chu, S., X. Xie, C. Payan and U. Stochaj, 2023 Valosin containing protein (VCP): initiator, modifier, and potential drug target for neurodegenerative diseases. Mol Neurodegener 18: 52.

Cleveland, D. W., and J. D. Rothstein, 2001 From charcot to lou gehrig: deciphering selective motor neuron death in als. Nature Reviews Neuroscience 2: 806–819.

Colombo, S. F., and E. Fasana, 2011 Mechanisms of insertion of tail-anchored proteins into the membrane of the endoplasmic reticulum. Curr Protein Pept Sci 12: 736–742.

Deivasigamani, S., H. K. Verma, R. Ueda, A. Ratnaparkhi and G. S. Ratnaparkhi, 2014 A genetic screen identifies Tor as an interactor of VAPB in a Drosophila model of amyotrophic lateral sclerosis. Biology Open 3: 1127–1138.

Doble, A., 1996 The pharmacology and mechanism of action of riluzole. Neurology 47: S233–241.

Ewens, C. A., S. Panico, P. Kloppsteck, C. McKeown, I. O. Ebong et al., 2014 The p97-FAF1 protein complex reveals a common mode of p97 adaptor binding. J Biol Chem 289: 12077–12084.

Fasana, E., M. Fossati, A. Ruggiano, S. Brambillasca, C. C. Hoogenraad et al., 2010 A VAPB mutant linked to amyotrophic lateral sclerosis generates a novel form of organized smooth endoplasmic reticulum. FASEB J 24: 1419–1430.

Genevini, P., G. Papiani, A. Ruggiano, L. Cantoni, F. Navone et al., 2014 Amyotrophic lateral sclerosis-linked mutant VAPB inclusions do not interfere with protein degradation pathways or intracellular transport in a cultured cell model. PLoS One 9: e113416.

Griciuc, A., L. Aron, M. J. Roux, R. Klein, A. Giangrande et al., 2010 Inactivation of VCP/ter94 suppresses retinal pathology caused by misfolded rhodopsin in Drosophila. PLoS Genet 6.

Haas, I. G., 1994 BiP (GRP78), an essential hsp70 resident protein in the endoplasmic reticulum. Experientia 50: 1012–1020.

Han, S. M., H. Tsuda, Y. Yang, J. Vibbert, P. Cottee et al., 2012 Secreted VAPB/ALS8 major sperm protein domains modulate mitochondrial localization and morphology via growth cone guidance receptors. Dev Cell 22: 348–362.

Hegde, R. S., and R. J. Keenan, 2011 Tail-anchored membrane protein insertion into the endoplasmic reticulum. Nat Rev Mol Cell Biol 12: 787–798.

Hemion, C., J. Flammer and A. Neutzner, 2014 Quality control of oxidatively damaged mitochondrial proteins is mediated by p97 and the proteasome. Free Radic Biol Med 75: 121–128.

Heo, J. M., N. Livnat-Levanon, E. B. Taylor, K. T. Jones, N. Dephoure et al., 2010 A stress-responsive system for mitochondrial protein degradation. Mol Cell 40: 465–480.

Huttlin, E. L., L. Ting, R. J. Bruckner, F. Gebreab, M. P. Gygi et al., 2015 The BioPlex Network: A Systematic Exploration of the Human Interactome. Cell 162: 425–440.

Ibrahim, I. M., D. H. Abdelmalek and A. A. Elfiky, 2019 GRP78: A cell’s response to stress. Life Sci 226: 156–163.

James, C., and R. H. Kehlenbach, 2021 The Interactome of the VAP Family of Proteins: An Overview. Cells 10.

Johnson, J. O., J. Mandrioli, M. Benatar, Y. Abramzon, V. M. Van Deerlin et al., 2010 Exome sequencing reveals VCP mutations as a cause of familial ALS. Neuron 68: 857–864.

Ju, J. S., R. A. Fuentealba, S. E. Miller, E. Jackson, D. Piwnica-Worms et al., 2009 Valosin-containing protein (VCP) is required for autophagy and is disrupted in VCP disease. J Cell Biol 187: 875–888.

Ju, J. S., S. E. Miller, P. I. Hanson and C. C. Weihl, 2008 Impaired protein aggregate handling and clearance underlie the pathogenesis of p97/VCP-associated disease. J Biol Chem 283: 30289–30299.

Kaduskar, B., D. Trivedi and G. S. Ratnaparkhi, 2020 Caspar SUMOylation regulates Drosophila lifespan. MicroPubl Biol 2020.

Kanekura, K., I. Nishimoto, S. Aiso and M. Matsuoka, 2006 Characterization of amyotrophic lateral sclerosis-linked P56S mutation of vesicle-associated membrane protein-associated protein B (VAPB/ALS8). J Biol Chem 281: 30223–30233.

Kim, M., J. H. Lee, S. Y. Lee, E. Kim and J. Chung, 2006 Caspar, a suppressor of antibacterial immunity in Drosophila. Proceedings of the National Academy of Sciences of the United States of America 103: 16358–16363.

Kuijpers, M., V. van Dis, E. D. Haasdijk, M. Harterink, K. Vocking et al., 2013 Amyotrophic lateral sclerosis (ALS)-associated VAPB-P56S inclusions represent an ER quality control compartment. Acta Neuropathol Commun 1: 24.

Kurland, L. T., and D. W. Mulder, 1955 Epidemiologic investigations of amyotrophic lateral sclerosis. 2. Familial aggregations indicative of dominant inheritance. II. Neurology 5: 249–268.

Larroquette, F., L. Seto, P. L. Gaub, B. Kamal, D. Wallis et al., 2015 Vapb/Amyotrophic lateral sclerosis 8 knock-in mice display slowly progressive motor behavior defects accompanying ER stress and autophagic response. Hum Mol Genet 24: 6515–6529.

Leblond, C. S., H. M. Kaneb, P. A. Dion and G. A. Rouleau, 2014 Dissection of genetic factors associated with amyotrophic lateral sclerosis. Exp Neurol 262 Pt B: 91-101.

Lee, A. S., 2005 The ER chaperone and signaling regulator GRP78/BiP as a monitor of endoplasmic reticulum stress. Methods 35: 373–381.

Lee, J. J., J. K. Park, J. Jeong, H. Jeon, J. B. Yoon et al., 2013 Complex of Fas-associated factor 1 (FAF1) with valosin-containing protein (VCP)-Npl4-Ufd1 and polyubiquitinated proteins promotes endoplasmic reticulum-associated degradation (ERAD). J Biol Chem 288: 6998–7011.

Leon, A., and D. McKearin, 1999 Identification of TER94, an AAA ATPase protein, as a Bam-dependent component of the Drosophila fusome. Mol Biol Cell 10: 3825–3834.

Lim, J., and Z. Yue, 2015 Neuronal aggregates: formation, clearance, and spreading. Dev Cell 32: 491–501.

Loewen, C. J., and T. P. Levine, 2005 A highly conserved binding site in vesicle-associated membrane protein-associated protein (VAP) for the FFAT motif of lipid-binding proteins. J Biol Chem 280: 14097–14104.

Lord, J. M., A. Ceriotti and L. M. Roberts, 2002 ER dislocation: Cdc48p/p97 gets into the AAAct. Curr Biol 12: R182–184.

Meyer, H., M. Bug and S. Bremer, 2012 Emerging functions of the VCP/p97 AAA-ATPase in the ubiquitin system. Nat Cell Biol 14: 117–123.

Miller, T. M., M. E. Cudkowicz, A. Genge, P. J. Shaw, G. Sobue et al., 2022 Trial of Antisense Oligonucleotide Tofersen for SOD1 ALS. N Engl J Med 387: 1099–1110.

Mitchell, J. D., and G. D. Borasio, 2007 Amyotrophic lateral sclerosis. Lancet 369: 2031–2041.

Mitne-Neto, M., M. Machado-Costa, M. C. Marchetto, M. H. Bengtson, C. A. Joazeiro et al., 2011 Downregulation of VAPB expression in motor neurons derived from induced pluripotent stem cells of ALS8 patients. Hum Mol Genet 20: 3642–3652.

Mitne-Neto, M., C. R. Ramos, D. C. Pimenta, J. S. Luz, A. L. Nishimura et al., 2007 A mutation in human VAP-B--MSP domain, present in ALS patients, affects the interaction with other cellular proteins. Protein Expr Purif 55: 139–146.

Moustaqim-Barrette, A., Y. Q. Lin, S. Pradhan, G. G. Neely, H. J. Bellen et al., 2014 The amyotrophic lateral sclerosis 8 protein, VAP, is required for ER protein quality control. Hum Mol Genet 23: 1975–1989.

Mukkavalli, S., J. A. Klickstein, B. Ortiz, P. Juo and M. Raman, 2021 The p97-UBXN1 complex regulates aggresome formation. J Cell Sci 134.

Mulakkal, N. C., P. Nagy, S. Takats, R. Tusco, G. Juhasz et al., 2014 Autophagy in Drosophila: from historical studies to current knowledge. Biomed Res Int 2014: 273473.

Murphy, S. E., and T. P. Levine, 2016 VAP, a Versatile Access Point for the Endoplasmic Reticulum: Review and analysis of FFAT-like motifs in the VAPome. Biochim Biophys Acta 1861: 952–961.

Nalbandian, A., K. J. Llewellyn, M. Kitazawa, H. Z. Yin, M. Badadani et al., 2012 The homozygote VCP(R(1)(5)(5)H/R(1)(5)(5)H) mouse model exhibits accelerated human VCP-associated disease pathology. PLoS One 7: e46308.

Nishimura, A. L., M. Mitne-Neto, H. C. A. Silva, A. Richieri-Costa, S. Middleton et al., 2004 A Mutation in the Vesicle-Trafficking Protein VAPB Causes Late-Onset Spinal Muscular Atrophy and Amyotrophic Lateral Sclerosis. The American Journal of Human Genetics 75: 822–831.

Nishimura, Y., M. Hayashi, H. Inada and T. Tanaka, 1999 Molecular cloning and characterization of mammalian homologues of vesicle-associated membrane protein-associated (VAMP-associated) proteins. Biochem Biophys Res Commun 254: 21–26.

Paganoni, S., E. A. Macklin, S. Hendrix, J. D. Berry, M. A. Elliott et al., 2020 Trial of Sodium Phenylbutyrate-Taurursodiol for Amyotrophic Lateral Sclerosis. N Engl J Med 383: 919–930.

Papiani, G., A. Ruggiano, M. Fossati, A. Raimondi, G. Bertoni et al., 2012 Restructured endoplasmic reticulum generated by mutant amyotrophic lateral sclerosis-linked VAPB is cleared by the proteasome. J Cell Sci 125: 3601–3611.

Park, M. Y., J. H. Moon, K. S. Lee, H. I. Choi, J. Chung et al., 2007 FAF1 suppresses I kappa B kinase (IKK) activation by disrupting the IKK complex assembly. Journal of Biological Chemistry 282: 27572–27577.

Pasinelli, P., and R. H. Brown, 2006 Molecular biology of amyotrophic lateral sclerosis: insights from genetics. Nature Reviews Neuroscience 7: 710–723.

Pennetta, G., P. R. Hiesinger, R. Fabian-Fine, I. A. Meinertzhagen and H. J. Bellen, 2002 Drosophila VAP-33A directs bouton formation at neuromuscular junctions in a dosage-dependent manner. Neuron 35: 291–306.

Prause, J., A. Goswami, I. Katona, A. Roos, M. Schnizler et al., 2013 Altered localization, abnormal modification and loss of function of Sigma receptor-1 in amyotrophic lateral sclerosis. Hum Mol Genet 22: 1581–1600.

Rabinovich, E., A. Kerem, K. U. Frohlich, N. Diamant and S. Bar-Nun, 2002 AAA-ATPase p97/Cdc48p, a cytosolic chaperone required for endoplasmic reticulum-associated protein degradation. Mol Cell Biol 22: 626–634.

Rai, M., M. Curley, Z. Coleman, A. Nityanandam, J. Jiao et al., 2021 Analysis of proteostasis during aging with western blot of detergent-soluble and insoluble protein fractions. STAR Protoc 2: 100628.

Ratnaparkhi, A., G. M. Lawless, F. E. Schweizer, P. Golshani and G. R. Jackson, 2008 A Drosophila Model of ALS: Human ALS-Associated Mutation in VAP33A Suggests a Dominant Negative Mechanism. PLoS ONE 3: e2334.

Reis-Rodrigues, P., G. Czerwieniec, T. W. Peters, U. S. Evani, S. Alavez et al., 2012 Proteomic analysis of age-dependent changes in protein solubility identifies genes that modulate lifespan. Aging Cell 11: 120–127.

Renton, A. E., A. Chio and B. J. Traynor, 2014 State of play in amyotrophic lateral sclerosis genetics. Nat Neurosci 17: 17–23.

Ritson, G. P., S. K. Custer, B. D. Freibaum, J. B. Guinto, D. Geffel et al., 2010 TDP-43 Mediates Degeneration in a Novel Drosophila Model of Disease Caused by Mutations in VCP/p97. Journal of Neuroscience 30: 7729–7739.

Ross, C. A., and M. A. Poirier, 2004 Protein aggregation and neurodegenerative disease. Nat Med 10 Suppl: S10–17.

Ryu, S. W., and E. Kim, 2001 Apoptosis induced by human Fas-associated factor 1, hFAF1, requires its ubiquitin homologous domain, but not the Fas-binding domain. Biochem Biophys Res Commun 286: 1027–1032.

Ryu, S. W., S. J. Lee, M. Y. Park, J. Jun, Y. K. Jung et al., 2003 Fas-associated factor 1, FAF1, is a member of Fas death-inducing signaling complex. Journal of Biological Chemistry 278: 24003–24010.

Seguin, S. J., F. F. Morelli, J. Vinet, D. Amore, S. De Biasi et al., 2014 Inhibition of autophagy, lysosome and VCP function impairs stress granule assembly. Cell Death Differ 21: 1838–1851.

Skehel, P. A., K. C. Martin, E. R. Kandel and D. Bartsch, 1995 A VAMP-binding protein from Aplysia required for neurotransmitter release. Science 269: 1580–1583.

Suzuki, H., K. Kanekura, T. P. Levine, K. Kohno, V. M. Olkkonen et al., 2009 ALS-linked P56S-VAPB, an aggregated loss-of-function mutant of VAPB, predisposes motor neurons to ER stress-related death by inducing aggregation of co-expressed wild-type VAPB. J Neurochem 108: 973–985.

Szegezdi, E., A. Duffy, M. E. O’Mahoney, S. E. Logue, L. A. Mylotte et al., 2006 ER stress contributes to ischemia-induced cardiomyocyte apoptosis. Biochem Biophys Res Commun 349: 1406–1411.

Tendulkar, S., S. Hegde, L. Garg, A. Thulasidharan, B. Kaduskar et al., 2022 Caspar, an adapter for VAPB and TER94, modulates the progression of ALS8 by regulating IMD/NFkappaB-mediated glial inflammation in a Drosophila model of human disease. Hum Mol Genet 31: 2857–2875.

Teuling, E., S. Ahmed, E. Haasdijk, J. Demmers, M. O. Steinmetz et al., 2007 Motor Neuron Disease-Associated Mutant Vesicle-Associated Membrane Protein-Associated Protein (VAP) B Recruits Wild-Type VAPs into Endoplasmic Reticulum-Derived Tubular Aggregates. Journal of Neuroscience 27: 9801–9815.

Tresse, E., F. A. Salomons, J. Vesa, L. C. Bott, V. Kimonis et al., 2010 VCP/p97 is essential for maturation of ubiquitin-containing autophagosomes and this function is impaired by mutations that cause IBMPFD. Autophagy 6: 217–227.

Tsakiri, E. N., G. P. Sykiotis, I. S. Papassideri, V. G. Gorgoulis, D. Bohmann et al., 2013 Differential regulation of proteasome functionality in reproductive vs. somatic tissues of Drosophila during aging or oxidative stress. FASEB J 27: 2407–2420.

Tsuda, H., S. M. Han, Y. Yang, C. Tong, Y. Q. Lin et al., 2008 The Amyotrophic Lateral Sclerosis 8 Protein VAPB Is Cleaved, Secreted, and Acts as a Ligand for Eph Receptors. Cell 133: 963–977.

van Blitterswijk, M., M. A. van Es, E. A. Hennekam, D. Dooijes, W. van Rheenen et al., 2012 Evidence for an oligogenic basis of amyotrophic lateral sclerosis. Hum Mol Genet 21: 3776–3784.

van den Boom, J., and H. Meyer, 2018 VCP/p97-Mediated Unfolding as a Principle in Protein Homeostasis and Signaling. Molecular Cell 69: 182–194.

Voeltz, G. K., E. M. Sawyer, G. Hajnoczky and W. A. Prinz, 2024 Making the connection: How membrane contact sites have changed our view of organelle biology. Cell 187: 257–270.

Wang, B., B. A. Maxwell, J. H. Joo, Y. Gwon, J. Messing et al., 2019 ULK1 and ULK2 Regulate Stress Granule Disassembly Through Phosphorylation and Activation of VCP/p97. Mol Cell 74: 742–757 e748.

Weihl, C. C., S. Dalal, A. Pestronk and P. I. Hanson, 2006 Inclusion body myopathy-associated mutations in p97/VCP impair endoplasmic reticulum-associated degradation. Hum Mol Genet 15: 189–199.

Wilson, D. M., 3rd, M. R. Cookson, L. Van Den Bosch, H. Zetterberg, D. M. Holtzman et al., 2023 Hallmarks of neurodegenerative diseases. Cell 186: 693–714.

Yadav, S., R. Thakur, P. Georgiev, S. Deivasigamani, H. Krishnan et al., 2018 RDGBalpha localization and function at membrane contact sites is regulated by FFAT-VAP interactions. J Cell Sci 131: jcs207985.

Yamanaka, K., Y. Sasagawa and T. Ogura, 2012 Recent advances in p97/VCP/Cdc48 cellular functions. Biochim Biophys Acta 1823: 130–137.

Yoshino, H., and A. Kimura, 2006 Investigation of the therapeutic effects of edaravone, a free radical scavenger, on amyotrophic lateral sclerosis (Phase II study). Amyotroph Lateral Scler 7: 241–245.

Zhang, W., A. Colavita and J. K. Ngsee, 2017 Mitigating Motor Neuronal Loss in C. elegans Model of ALS8. Sci Rep 7: 11582.

Zhang, X. Y., L. Gui, X. Y. Zhang, S. L. Bulfer, V. Sanghez et al., 2015 Altered cofactor regulation with disease-associated p97/VCP mutations. Proceedings of the National Academy of Sciences of the United States of America 112: E1705–E1714.

Zhao, Y. G., N. Liu, G. Y. Miao, Y. Chen, H. Y. Zhao et al., 2018 The ER Contact Proteins VAPA/B Interact with Multiple Autophagy Proteins to Modulate Autophagosome Biogenesis. Current Biology 28: 1234-+.

Zhu, J., S. Pittman, D. Dhavale, R. French, J. N. Patterson et al., 2022 VCP suppresses proteopathic seeding in neurons. Mol Neurodegener 17: 30.

