## Supplementary material for "Age-dependent dynamics of neuronal VAPB^ALS^ inclusions in the adult brain": Suppl. Figure 1

**A**

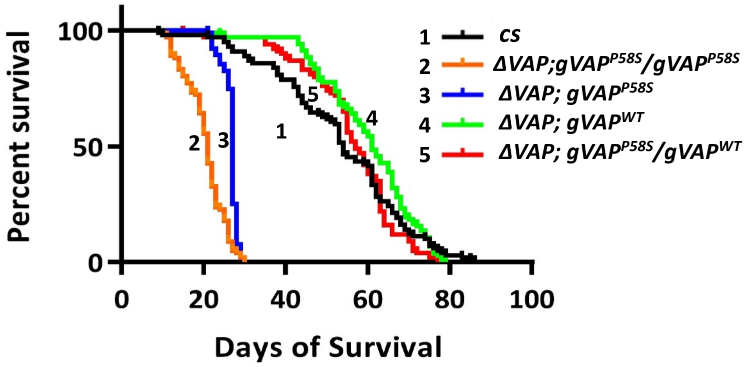


**B**

| | Genotypes | P-value, median lifespan |
|---|---|---|
| 1 | *+/+ (wild type)* | control, 54 |
| 2 | *ΔVAP; $gVAP^{P58S}/gVAP^{P58S}$* | ***, 21 |
| 3 | *ΔVAP; $gVAP^{P58S}$* | ***, 27 |
| 4 | *ΔVAP; $gVAP^{WT}$* | ns, 61 |
| 5 | *ΔVAP; $gVAP^{P58S}/gVAP^{WT}$* | ns, 57 |

**C**

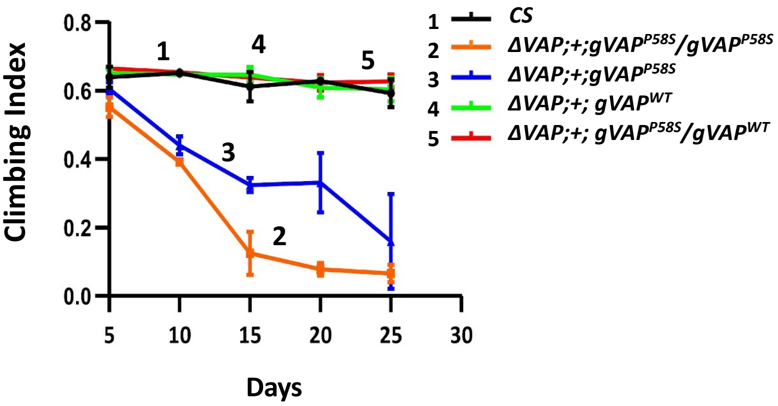


***Figure S1: ΔVAP;+; $gVAP^{P58S}$ (Null rescue) flies show lifespan defects and progressive motor degeneration***

**(A)** Survival plots for CS (wild type, master control, black curve, number 1), *ΔVAP;+; $gVAP^{P58S}/gVAP^{P58S}$* (orange curve, number 2), ΔVAP;+;$gVAP^{P58S}$ (blue curve, number 3), *ΔVAP;+;$gVAP^{WT}$* (light green curve, number 4) and *ΔVAP; $gVAP^{WT}/gVAP^{P58S}$* (red curve, number 5). Introduction of a *$gVAP^{wt}$* copy in the *ΔVAP* or *ΔVAP; $gVAP^{P58S}$* background rescues the lifespan defect. Curve comparison was done using log-rank (Mantel-Cox test). Combined p-value for the whole set is <0.001 (n=80-100).

**(B)** The table mentions the Median lifespan values for each set. Curve comparison was done using log-rank (Mantel-Cox test). p-values depict values of significance from log-rank test. The combined p-value for the whole set is <0.001 ( n=80-100 flies for each genotype.)

**(C)** Climbing index of *ΔVAP;+;$gVAP^{P58S}$* flies (blue) is significantly reduced as compared to wild-type control flies (black) when observed from day 5 to day 30 of the life of the flies. We observe a rescue in this decline upon adding a *$VAP^{WT}$* allele to the *$VAP^{P58S}$* genetic background (red). n=15-20 flies, N=1. Statistical analysis was done using an unpaired Student's t-test. The combined p-value for the whole set is <0.01
